# Proteome overabundance enables respiration but limitation onsets carbon overflow

**DOI:** 10.1101/2020.02.20.957662

**Authors:** Rahul Kumar, Aleksandr Illarionov, Petri-Jaan Lahtvee

**Affiliations:** Institute of Technology, University of Tartu, Estonia; Department of Chemistry and Biotechnology, Tallinn University of Technology, Estonia

**Keywords:** nutrient, mitochondria, metabolism, carbon overflow, ribosome, proteome glucose, nitrogen limitation, respiration, fermentation, yeast, cancer, biotechnology

## Abstract

Central carbon metabolism produces energy and precursor metabolites for biomass in heterotrophs. Carbon overflow yields metabolic byproducts and, here, we examined its dependency on nutrient and growth using the unicellular eukaryotic model organism *Saccharomyces cerevisiae*. We performed quantitative proteomics analysis together with metabolic modeling and found that proteome overabundance enabled respiration, and variation in energy efficiency caused distinct composition of biomass at different carbon to nitrogen ratio and growth rate. Our results showed that ceullar resource allocation for ribosomes was determinative of growth rate, but energy constrains on protein synthesis incepted carbon overflow by prioritizing abundance of ribosomes and glycolysis over mitochondria. We proved that glycolytic efficiency affected energy metabolism by making a trade-off between low and high energy production pathways. Finally, we summarized cellular energy budget underlying nutrient-responsive and growth rate-dependent carbon overflow, and suggested implications of results for bioprocesses and pathways relevant in cancer metabolism in humans.

## Introduction

Nutrient sensing and signaling is essential for proliferation and differentiation of cells. In prokaryotes and unicellular eukaryotes, nutrient act not only as substrate but also as signal for control of proliferation which requires an appreciation of the role of nutrient as signaling molecule and metabolite (Broach, 2012). Our study focuses on unicellular eukaryote, budding yeast *Saccharomyces cerevisiae*, which is considered a model organism for investigating eukaryal regulations as well as an industrial workhorse in biotechnology. In particular, we focus on the role of macronutrient carbon and nitrogen in carbon overflow metabolism in yeast. Carbon overflow is a metabolic response to diverse stimuli and, in the most prominent example, is described by the Warburg effect where respiring healthy mammalian cells shift metabolism to fermentation in cancer cells (Warburg, 1956). However, a different mechanism results carbon overflow in the Crabtree effect wherein presence of excess glucose represses respiration allowing aerobic glycolysis to be the main source of energy supply in several species of yeast (Crabtree, 1928; De Deken, 1966). Besides biomass and carbon dioxide (CO_2_) carbon overflow result in formation of metabolic byproducts such as organic acids in bacteria, ethanol in yeast and lactate in cancer cells (Vander Heiden et al., 2009). From a biosustainability viewpoint, investigating the role of macronutrients is important because e.g., nitrogen limitation is implicated in lipid metabolism that provides precursors for the production of oleochemicals such as biofuels (Yu et al., 2018).

Carbon overflow can be onset by a diverse set of stimuli in cells. A common reason for its onset is fast growth rate (e.g., bacteria and yeasts), but it can also be induced in slow growing cells (e.g., healthy human cells) confronting adverse conditions such as defect in nutrient sensing and signaling pathways, genetic or epigenetic modifications and environmental stress (Kumar et al., 2014; Lahtvee et al., 2016; Torrence and Manning, 2018). Carbon overflow shows a pathway preference for production of the cellular energy adenosine triphosphate (ATP) from glycolysis, a substrate level phosphorylation pathway with low ATP yield, over the electron transport chain (ETC), an oxidative phosphorylation pathway with high ATP yield (Vander Heiden et al., 2009). Together glycolysis, the tricarboxylic acid (TCA) cycle and the ETC constitute so called energy metabolism that is evolutionary conserved across organisms (Chen and Nielsen, 2019). The total ATP generated in energy metabolism is spent either on growth-associated energy costs (GAEC) or on energetic costs of non-growth associated maintenance (NGAM) (Chen and Nielsen, 2019). In a fully respiratory metabolism the ATP produced by consuming a catabolic substrate is mostly coupled to anabolic processes enabling growth by formation of biomass (GAEC) and relatively less energy expenditure occurs on NGAM as compared to fermentation (Chen and Nielsen, 2019; Molenaar et al., 2009; Shimizu and Matsuoka, 2019).

Glucose is the most common carbon source for *S. cerevisiae* and its biochemical breakdown produces ATP in glycolysis, a key biochemical pathway in the central carbon metabolism (CCM) that is involved both in energy generation but also provides precursor molecules for formation of biomass. In yeast, an uncoupling of ATP supply-demand for synthesis of biomass can occur over a range of nutrient and growth conditions causing respirofementative metabolism and formation of byproducts (Larsson et al., 1993). In our previous studies, we showed that at the same growth rate (i.e., at physiological steady state), changes in carbon to nitrogen ratio (C:N ratio) can shift metabolism from respiration to fermentation in bacteria and yeast (Kumar and Shimizu, 2010; Zhang et al., 2011). A physiological steady state for suspension cells is obtained by using chemostats where constant parameters can be maintained during cultivation (Kumar and Shimizu, 2011). Such steady states provide a model system for studying cellular regulation and, previously, allowed us to map interaction of evolutionary conserved nutrient-responsive pathways, that are also implicated in cancer, namely, sucrose fermenting type 1 (Snf1), an AMP-activated kinase (AMPK), and the target of rapamycin complex (TORC1) in *S. cerevisiae* (Zhang et al., 2011).

In recent reports one of differences in respiration and growth rate-dependent carbon overflow is attributed to cellular resource allocation strategy in bacteria and yeast (Basan et al., 2015; Metzl-Raz et al., 2017; Peebo et al., 2015). Since our previous studies indicate a crucial role of nutrient in the onset of carbon overflow, we asked whether different resource allocation strategy might also be important at the same growth rate when changes in cellular environment shift metabolism from respiration to fermentation (Kumar and Shimizu, 2011; Zhang et al., 2011). Further, a comparison of carbon overflow at the same growth rate caused in response to changes in cellular environment (nutrient-responsive) with one caused by changes in growth rate (growth rate-dependent) can allow us to uncover underlying diversity in metabolism as well identify potential mechanisms that control energy budget in yeast. In essence, we focused on the role of nutrient in relation to carbon overflow at different growth rates due to its relevance in biotechnology applications and fundamental importance in metabolic disorders, such as cancer and diabetes.

We performed quantitative proteomics analysis together with metabolic modeling approach using data from the physiological steady states of *S. cerevisiae* cultures. Briefly, our results showed that proteome overabundance enabled respiration, and variation in energy efficiency caused distinct composition of biomass at different carbon to nitrogen ratio and growth rate. Our results showed that cellular resource allocation for ribosomes was determinative of growth rate, but it was energy constrains on protein synthesis that led to onset of carbon overflow by prioritizing abundances of ribosome and glycolysis over mitochondria. We proved that glycolytic flux impinged on energy metabolism by making trade-off between high and low energy yield pathways in the buddying yeast. Finally, we summarized cellular energy budget underlying diversity of metabolism in the both nutrient-responsive and growth rate-dependent carbon overflows in comparison to fully respiring conditions in *S. cerevisiae*, and suggested practical implications of our results in one carbon metabolism, aspartate biosynthesis and fatty acid biosynthesis for bioprocesses and pathways in cancer.

## Results

### Diversity in carbon overflow: nutrient-responsive vs growth rate-dependent

We used the budding yeast *S. cerevisiae* CEN.PK 113-7D, a prototypic strain, to conduct 21 steady state chemostats at a dilution rate (D) of 0.1 h^−1^ (Figure 1, S1A). The experiments were performed in biological triplicates under seven different step-wise nitrogen gradients and thus changing the culture environment from nitrogen excess (C:N ratio, 4) to limitation (C:N ratio, 75) while maintaining a constant glucose concentration (10 g/l) (Figure 1A, S1A). This allowed us to identify the critical C:N ratio of 22 and a further increase in C:N ratio resulted in a nutrient-responsive shift of metabolism from respiration to fermentation at the same growth rate (D, 0.1 h^−1^) (Figure 1B, S1A). The metabolic shift at slow growth (D, 0.1 h^−1^) resulted in production of metabolic byproducts at C:N ratio (≥22) (Figure 1B, S1A). To compare diverse carbon overflow, i.e., nutrient-responsive vs growth rate-dependent, we conducted four independent steady state experiments: two at D = 0.26 h^−1^ and two at D = 0.32 h^−1^ (further referred to as fast growth) using the reference C:N ratio (4) (Figure 1A, S1A). The fast growth rate were chosen based on previous studies characterizing carbon overflow conditions (Larsson et al., 1993). The dilution rate 0.1 h^−1^ resulted in a slow growth representing only about 25% of maximum specific growth rate, while fast growth (0.26 h^−1^ and 0.32 h^−1^) reached up to 65-80% of maximum specific growth rate (approximately 0.4 h^−1^) achieved using the same minimal medium and cultivation conditions for *S. cerevisiae* (Canelas et al., 2010). The data from slow growth (C:N ratio 4 to 10) and fast-growth (both dilution rates) conditions were separately combined due to their similarities in physiology (Figure 1B-D, S1A). Both, slow growth (C:N ratio ≥22) and fast growth rates (C:N ratio 4) showed a decrease in biomass yield that was accompanied by formation of byproducts (e.g., ethanol, acetate) and a metabolic shift from respiration to fermentation was indicated by the respiratory quotient (RQ >1) (Figure 1B-D). The proteome yield (protein (g)/ dry cellular weight (dcw) (g)) at the reference condition (D, 0.1 h^−1^; C:N ratio 4) was 0.50±0.08 g/g (50%), but yield was reduced to 0.27±0.03 g/g (27%) at the highest C:N ratio (75) (Figure 1C). The critical C:N ratio (22) showed 20% less biomass, but a similar proteome yield (g/g) as the reference (Figure 1C). Moreover, no detectable ethanol concentration was detected and a fully respiratory metabolism was maintained (RQ 1) at the critical C:N ratio (Figure 1B, 1D). At fast growth (C:N ratio 4) biomass yield was reduced by about 20% and unlike the critical C:N ratio (22) at slow growth here was 18% decrease (Student’s t-test, 0.0001) in proteome content i.e., 0.41±0.05 g/g dcw (41%) compared to the reference that showed a respirofermentative metabolism (RQ >1) (Figure 1C-D, S1A). A reduction in biomass but not in proteome as well lack of ethanol prompted us to examine the total carbon balance in all our experiments (S1A). We found that at the critical C:N ratio (22) condition approximately 9% carbon was missing in the measured fluxes at the slow growth (D, 0.1 h^−1^). To understand distribution of this missing carbon the flux balance analysis (FBA) was performed that predicted distribution of extracellular carbon fluxes over several metabolic byproducts, namely formate (C1), glycerol, pyruvate (C3), and 2-oxaloacetate (C5) (Figure 1B, S1A, S1B). Among these byproducts, formate was suggested with the highest concentration probability (S1B). Because of this, we re-analyzed our extracellular metabolome samples using formate in the standards and detected production of formate (C1) at the critical and higher C:N ratios (≥22) at the slow growth rate conditions thereby validating the model prediction (Figure 1B, S1A, S1B). We did not find formate either at the slow growth (C:N ratio 4-10) or at the fast growth (C:N ratio 4) (Figure 1B, S1A). This indicates that at the slow growth (D, 0.1 h^−1^) the critical C:N ratio (22) provides a poised nutrient status as at the higher C:N ratios (≥22) nitrogen was insufficient to consume all available glucose as indicated by the residual glucose in the culture environment (Figure 1D, S1A). We determined that a specific glucose uptake rate (mmol/dcw(g)/h) higher than 2.0±0.3 incepted respirofermentative metabolism while at a lower rate a fully respiratory metabolism was maintained (Figure 1B-D, S1A). The cut-off specific glucose uptake rate value for carbon overflow was consistent with our previous study where respirofermentative metabolism was induced by environmental stress (Lahtvee et al., 2016). Overall, both nutrient-responsive and growth rate-dependent carbon overflow exhibit a limitation of available proteome, as indicated by reduced protein yields, while sustaining respirofermentative metabolism (Figure 1C).

**Figure 1:**
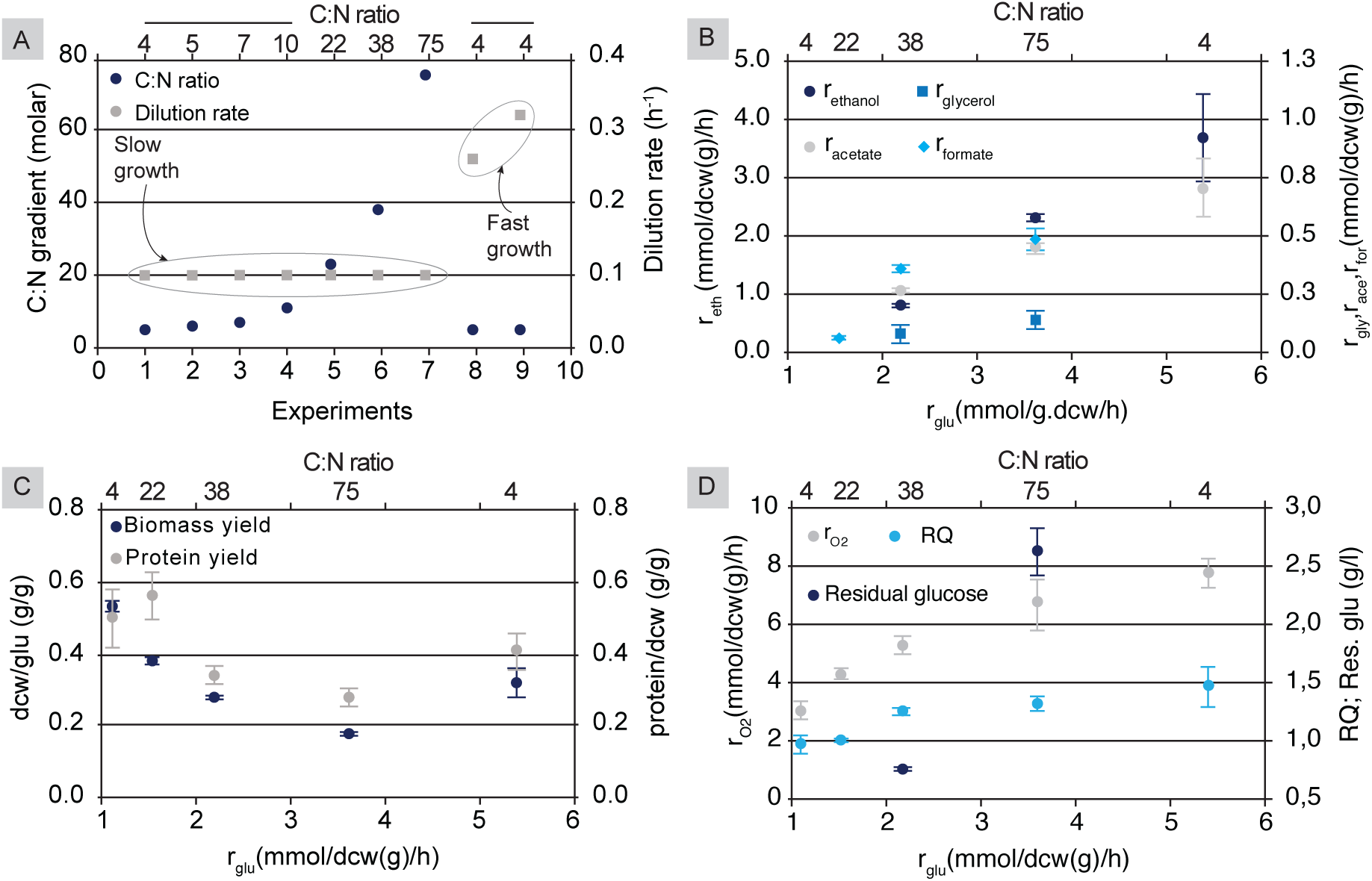
Diversity in carbon overflow: nutrient-responsive vs growth rate-dependent. (A) Schematic of chemostat experiments. Primary x- axis represents number of conditions. Primary y- axis shows glucose to ammonium sulfate molar ratio (C:N ratio). C:N ratio (molar) for each condition is represented on secondary x- axis. Chemostat dilution rate (h^−1^) is shown on secondary y- axis. The dilution rate 0.1 h^−1^ represents slow growth and indicated by a circle. The dilution rate 0.26 and 0.32 h^−1^ together indicate fast growth as marked by the circle. Each slow growth is experiment is conducted in biological triplicate and the fast growth is average of four experiments (two biological duplicates at 0.26 h^−1^ and two at 0.32 h^−1^). (B) Measured extra-cellular fluxes showing specific metabolite production rate (mmol/dcw(g)/h). Primary x- axis shows specific glucose uptake rate (r_glu_), primary y- axis shows specific ethanol production rate (r_eth_), secondary y- axis shows specific glycerol, acetate and formate production rates, respectively (r_gly,_ r_ace_, and r_for_). C:N ratio (molar) for each condition is indicated on secondary x- axis. (C) Biomass yield on glucose (dcw(g)/glu(g)) and protein yield in biomass (protein (g)/biomass (g)) is plotted on primary y- axis and secondary y- axis, respectively, with respect to specific glucose uptake rate on primary x- axis. (D) Specific oxygen uptake rate (r_O2_) on primary y- axis and respiratory quotient on secondary y- axis are plotted as a function of specific glucose uptake rate (r_glu_) on primary x- axis. Respiratory quotient (RQ). Residual glucose (g/l) is indicated for C:N ratio 38 and 75 on secondary y- axis. Data are mean ±SD. See also supplementary S1A and S1B.

### Distinct proteome profiles underline diversity in carbon overflow

We selected four different C:N ratio conditions (4, 22, 38, 75) in biological triplicate at the slow growth and three biological triplicate samples at the fast growth rate (C:N ratio 4) for a quantitative proteome analysis (Figure 2A). Due to the high similarity in physiology data with C:N ratio 4, we did not consider samples from C:N ratio 5-10 for the proteome analysis (S1A). We focused on five distinct physiology conditions, namely, reference (C:N ratio 4), the critical C:N ratio (22), nutrient-responsive carbon overflow (C:N ratios 38 and 75) at slow growth rate and growth rate-dependent carbon overflow (C:N ratio 4) at fast growth rate (Figure 2A, 2B). Further, in pairwise analysis, we selected C:N ratio (38) as representative condition for a nutrient-responsive carbon overflow as changes in C:N ratio (75) samples were similar but often showed a greater magnitude of change, and our focus was on identifying changes immediately adjacent to the critical C:N ratio that potentially onset nutrient-responsive carbon overflow (Figure 2C-D, S2A, S2B).

**Figure 2:**
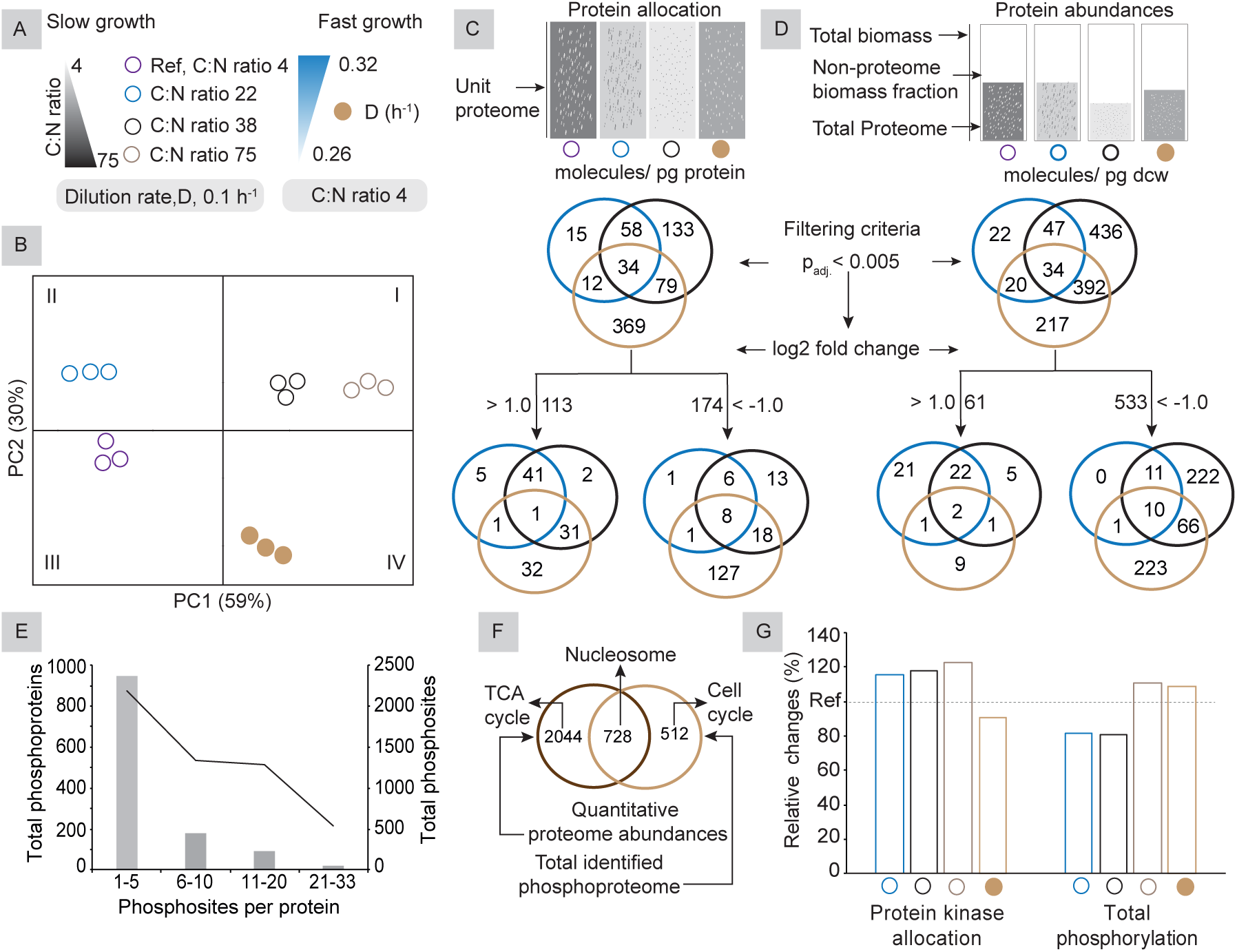
Distinct proteome profiles underline diversity in carbon overflow. (A) Describes design elements of the figure that are used throughout this manuscript where open circles with distinct colors indicate different C:N ratio at slow growth and filled circle indicates fast growth at the reference C:N ratio. (B) Principal component analysis (PCA) plot of quantitative proteomics data from biological triplicate experiments. The PC1 covers 59% of the variance in proteome abundances and separates experiments based on a fully respiratory or respirofementative metabolism. The PC2 covers additional 30% of the variance separating conditions based on growth rate. Since the C:N ratio 38 and 75 indicate biological similarity the C:N ratio 38 is shown as a representative of the both for visual clarity in some of the figures. (C) Schematic of comparison of protein molecules in the unit amount of proteome (molecules / picogram protein) represents protein allocation across the conditions. The first Venn diagram indicates significant proteins for each condition relative to the reference based on adjusted p- value (p_adj._ <0.005). The second Venn diagrams shows proteins with log2 fold changes (> 1 or < -1) within the significant proteome (p_adj._ <0.005) as considered in the first Venn diagram. See also supplementary S2A. (D) Schematic of comparison of protein molecules in the unit amount of biomass (measured as dry cellular weight, dcw) (molecules / picogram dcw) describes the protein abundances across the conditions. The same filtering criteria as described above (2C) is used. See also supplementary S2B. (E) Total phosphoproteome detected is categorized based on number of phosphosites per protein. Only phosphoproteins with more than 2 peptides in the MS data were considered in the analysis. The primary x- axis shows number of phosphosites per protein indicating most phosphoproteins in *S. cerevisiae* contain 1-5 phosphorylation modifications. The primary y- axis represents bars indicates total number of phosphoproteins while secondary y- axis represents line indicating total number of phosphosites. Bar represents primary y- axis and line indicates secondary y- axis. See also supplementary S2C. (F) The Venn diagram represents an evaluation of total estimated quantitative proteome and phosphoproteome and shows the coverage of phosphorylation in the proteome of *S. cerevisiae*. See also supplementary S2D. (G) Analysis of relative changes in proteome allocation to kinases, phosphorylating enzyme proteins, based on quantitative proteomics data (S2A) and relative changes in total phosphorylation based on intensities in phosphoproteome data (S2C).

A systems level analysis of absolute quantitative proteome abundances was performed by determining variances in our data using previously reported method (Lahtvee et al., 2017). The results in principal component analysis (PCA) showed most of variance (59%) on principal component (PC) 1 allowing a separation based on respiration (reference, C:N ratio of 22) or fermentation (fast growth, C:N ratio of 38 and 75) (Figure 2B). Additional variance (30%) in data on PC2 distinguished experiments mainly based on growth rate (Figure 2B). We used proteome-normalized data (protein molecules/ picogram proteome) to understand allocation differences and biomass-normalized data (molecules / picogram dcw) to understand abundance changes in each condition (Figure 2C-D, S2A, S2B). In both pairwise comparisons, the same filtering criteria were used to determine significant protein differences relative to the reference i.e., first, based only on the adjusted p-value (p_adj._ <0.005), and, second, based on a combination of the adjusted p-value (p_adj._ <0.005) and log2 fold change (log_2_FC >1 or < -1.0) (Figure 2C-D, S2A, S2B). The allocation analysis showed significant (p_adj._ <0.005) changes for 700 proteins or 25% of total identified proteins (Figure 2C, S2A). The abundance analysis suggested significant changes (p_adj._ <0.005) for 1168 proteins or 42% of total identified proteins (Figure 2D, S2B). In the both analyses more proteins showed a decrease (p_adj._ <0.005, log_2_ FC < -1.0) than an increase (p_adj._ <0.005, log_2_ FC >1.0) suggesting a general overabundance of proteome at the slow growth reference condition (Figure 2C-D, S2A, S2B). Among all the conditions least changes for allocation or abundances were found at the critical C:N ratio (22) that showed a respiratory metabolism similar to the reference (Figure 1B, 2C-D). In the instances of carbon overflow a contrast was noticed between proteins allocation and abundance changes between C:N ratio (38) vs fast growth (4) indicating that these conditions might implicate different mechanisms in the onset of overflow metabolism (Figure 2C-D). In allocation analysis, the nutrient-responsive carbon overflow showed less protein changes (p_adj._ <0.005) than the growth rate-dependent carbon overflow while reverse was the case for in abundance analysis (Figure 2C-D). This contrast was a reflection of protein yield (g/g) differences between nutrient-responsive and growth rate-dependent-carbon overflow as compared with the reference (Figure 1C, 2C-D).

As a significant part of proteome is considered regulated by post-translational modifications (PTMs), the most prominently by phosphorylation (Vlastaridis et al., 2017), and therefore we performed a relative phosphoproteome analysis using the same samples as in the quantitative abundance analysis (Figure 2E-G, S2C). We identified a total of 5348 phosphosites in 1242 different proteins confirming previous reports of the presence of large-scale phosphorylation modification in *S. cerevisiae* (Figure 2E, S2C) (Vlastaridis et al., 2017). Most of the phosphoproteome contained 1-5 phosphosites per protein and only a very few proteins were associated with more than 20 phosphosites in yeast (Figure 2E, S2C). A comparative analysis of absolute quantitative proteome and phosphoproteome showed that nearly 60% of phosphoproteins were also detected in our abundance data (Figure 2F). Further, we performed a gene-set enrichment analysis using quantitative proteome and phosphoproteome data, and identified the significant GO terms (Figure 2F, S2D). In analysis of relative changes in proteome allocation to kinases, phosphorylating enzyme proteins, based on quantitative proteomics data, and relative changes in total phosphorylation, based on intensities in phosphoproteome data, we found that kinase-phosphorylation correlation was present only for the C:N ratio of 75 at slow growth, but not for other conditions in our experiments (Figure 2G, S2A, S2C). It suggested that global phosphorylation levels were not likely determinants for onset of carbon overflow but, as previously reported, individual enzyme phosphorylation events might be pertinent for such regulation (Oliveira and Sauer, 2012).

In the main, our systems level analysis of quantitative proteome data showed that a fully respiratory metabolism was characterized by protein overabundances. In the next section, we focus on functional analysis of proteome allocation (molecules/ pg protein) and follow it by functional abundance analysis (molecules/ pg dcw) in the later section.

### Proteome allocation to ribosomes is determinative of growth rate

In the previous section, we showed that the two types of carbon overflow exhibit distinct proteome profiles (Figure 2). To understand the functional significance of this distinction, we categorized the proteome allocation (molecules /pg protein) analysis data into the GO terms, namely, amino acid biosynthesis, glycolysis, ribosome and mitochondrion (Figure 3A, S2A). The proteome allocation for these GO terms covered more than 65% of quantified proteins (Figure 3A). The large-scale proteome allocation differences were driven by ribosomes and pertained to the growth rates (Figure 3A). However, at the same growth rate proteome allocation to the GO terms remained almost steady even as proteome yield (g/g) decreased up to 40% compared to the reference condition in the instances of nutrient-responsive carbon overflow (Figure 3A-B). Thus, proteome allocation differences were mostly implicated in the growth rate-dependent carbon overflow but the decrease in protein abundances was important in the nutrient-responsive carbon overflow (Figure 3A-B). The proteome allocation to the GO terms glycolysis: mitochondrion: ribosome showed a ratio of 1: 1.6: 1.4 at the slow growth but changed to 1: 2: 2.7 at fast growth rate (Figure 3A). The increase in the specific glucose uptake rates was independent of proteome allocation for glycolysis as the allocation remained nearly constant despite a significant reduction in protein yield (g/g) at higher C:N ratio (>22), suggesting it to be maximum possible allocation for glycolysis, at the slow growth rate and was significantly reduced at fast growth-rate (Figure 3C). Thus, the increase in the specific glucose uptake rate occurred despite reduced proteome allocation and decreased total protein yield indicating control of glycolysis was not solely dependent on allocation or abundances but on some other constrains and as has been previously reported implicates posttranscriptional regulation (Figure 3A-C) (Daran-Lapujade et al., 2007). Ribosomes reflected translation capacity for protein synthesis but an increase in their allocation at the faster growth did not lead to a similar increase in mitochondrion that would have been required for the ATP generation through respiration (Figure 3A). The increase in translation capacity was instead utilized for almost 3-fold increase in the specific growth rate compared with the reference by likely generating additional required ATP through the glycolysis resulting in carbon overflow and reduced protein yield (Figure 3A-C). At higher C:N ratio (>22), although total protein amount decreased up to 40% compared to the reference conditions, proteins allocated for ribosomes reduced only by few percentage points, resulting in slightly lower translation capacity (Figure 3A-C). At the fast growth rate an increase in the translation capacity was mainly directed towards maintaining growth as proteome allocation for both ATP generating pathways, namely glycolysis and mitochondrion was reduced significantly causing reduced protein yield and incepting carbon overflow (Figure 3A-C, 1B). The inception of the both nutrient-responsive and growth rate-dependent carbon overflow was marked by a reduction in proteome allocation to mitochondrion (Figure 3A-C). A further analysis of mitochondrion proteome allocation changes, using the child GO terms, showed that allocation for mitochondrial translation reduced nearly 40% under nitrogen limitation at slow growth but increased over 40% at the fast growth compared with the reference implicating it in growth (Figure 3C-D). The proteome allocation to the GO terms Fe-S cluster binding, vacuole (and its child terms), nitrogen metabolism and fatty biosynthesis increased in response to nitrogen limitation (Figure 3E-F). The increase in proteome allocation to the vacuole GO term (and associated child terms) suggested its potential role in replenishing nitrogen by protein turnover under nitrogen limitation conditions and is consistent with its previously reported function in yeast (Martin-Perez and Villen, 2017). The proteome allocation to GO terms, namely vacuole, nitrogen metabolism and fatty acid biosynthesis but not for except Fe-S culture binding reduced more than 50% under nitrogen excess at the fast growth rate (Figure 3E-F). These results indicate that nitrogen limitation and slow growth could be beneficial for production of fatty acids derived biochemicals, e.g., biofuels (Buijs et al., 2015).

**Figure 3:**
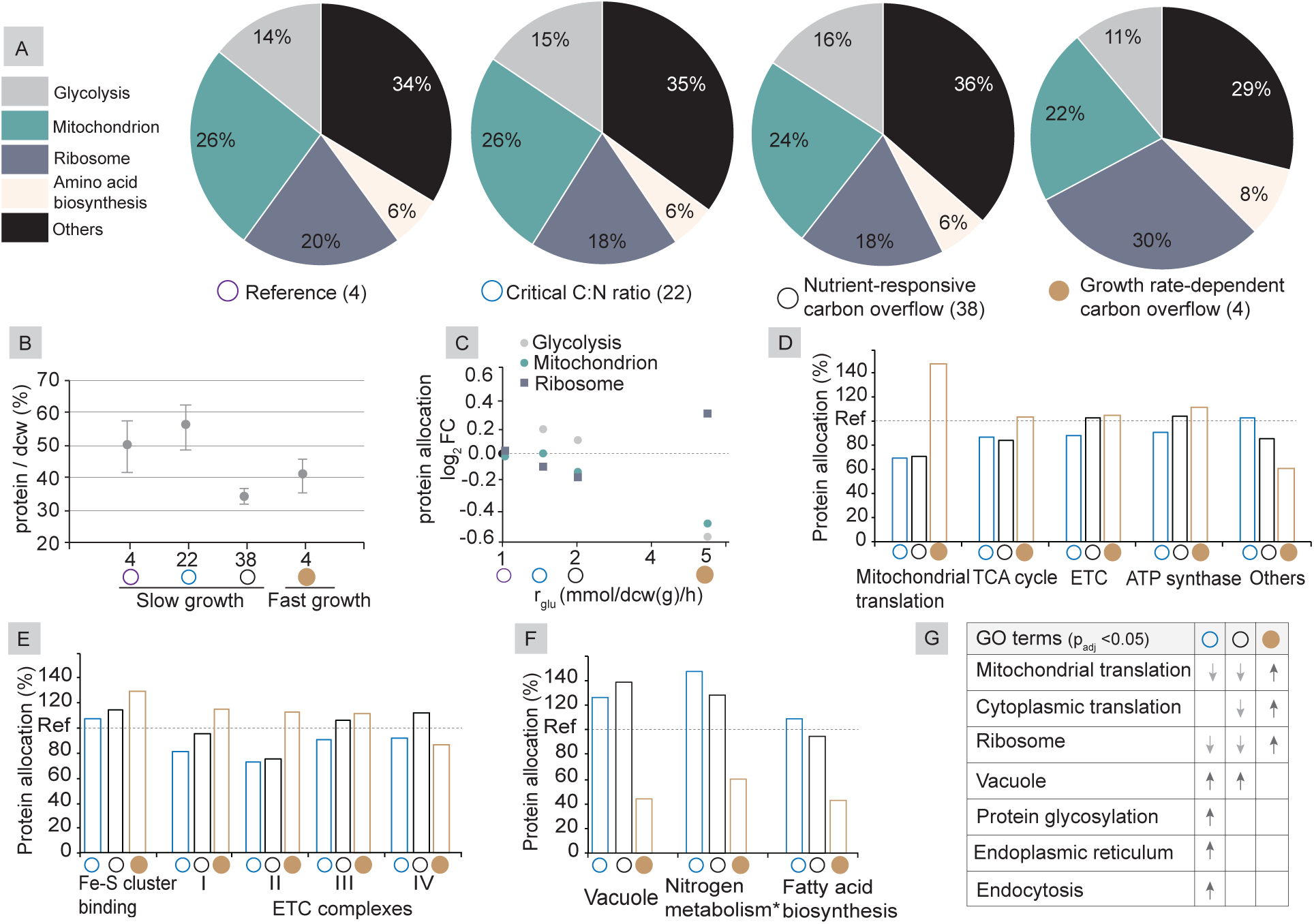
Proteome allocation to ribosomes is determinative of growth rate, but distinct translation constrains control diversity in carbon overflow. (A) Cellular resource allocation is indicated based on proteome normalized data (molecules / pg protein). The GO terms amino acid biosynthesis (GO:0008652), glycolysis (GO:0006096), ribosome (GO:0005840), and mitochondrion (GO:0005739) are accessed through the yeast genome database (yeastgenome.org) and cover over 65% of total estimated proteome. Each condition is illustrated by small empty or filled circle and described underneath the pie charts. (B) Percent protein yield in biomass across the conditions. Data are mean ±SD. Each condition is illustrated by small empty or filled circle. See also supplementary S1A. (C) Relative log_2_ fold changes in protein allocation (y- axis) for the GO terms glycolysis (GO:0006096), ribosome (GO:0005840), and mitochondrion (GO:0005739) compared with the reference plotted against the specific glucose uptake rate (mmol/dcw(g)/h) on the x-axis. Each condition is illustrated by small empty or filled circle. (D) Relative proteome allocation (%) to the mitochondrion child GO terms: mitochondrial translation (GO:0032543), the TCA cycle (GO:0006099), the electron transfer chain (ETC)* (constituted by the child GO terms – complex I-IV in 3E), ATP synthase (GO:0015986) and others indicate relative changes in the remainder of mitochondrial proteome. Each condition is illustrated by small empty or filled circle. (E) Relative proteome allocation (%) to the GO terms: iron-sulfur (Fe-S) cluster binding (GO:0051536), complex I - GO:0005747, complex II- GO:0005749, complex III- GO:0005750, complex IV - GO:0005751. Each condition is illustrated by small empty or filled circle. (F) Relative proteome allocation (%) to some of the GO terms in the proteome fraction categorized as “others” in 3A: vacuole (GO:0005773), nitrogen metabolism* -(GO:0006537, GO:0006542, GO:0015696, GO:0019740, GO:0006807) and fatty biosynthesis* (GO:0006631, GO:0006633). Each condition is illustrated by small empty or filled circle. (G) Gene-set analysis using proteome normalized data (S2A) shows significant GO terms (p_adj._<0.05). Direction of arrows indicate relative up or down regulation for a particular GO term relative to the reference. Each condition is illustrated by small empty or filled circle. ** See also supplementary S3. *The GO terms manually curated using the child GO terms. **Includes detailed list and also significant GO terms and TF analysis from the gene-set analysis.

### Distinct translation constrains control diversity in carbon overflow

Further, to statistically demonstrate significant differences in proteome allocation we performed gene-set analysis that has been previously reported (Varemo et al., 2013). This allowed us to identify significant (p_adj_ < 0.05) GO terms and TFs, including their directionalities (Figure 3G, S3). These results confirmed the discussion in previous paragraph but provided a few additional insights concerning protein translation and uniqueness of the critical C:N ratio proteome (Figure 3G, S3). The allocation differences in ribosomes appeared determinative for growth rate (Figure 3G). However, an absence of significant decrease in cytoplasmic translation and corresponding regulator TF Ifh11, a coactivator that regulates transcription of ribosomal protein (RP) genes, distinguished the critical C:N ratio from nutrient-responsive carbon overflow at the slow growth and the latter from the growth rate-dependent carbon overflow (Figure 3G, S3). The increase in proteome allocation to vacuole, under nitrogen limitation, was accompanied by the increase in TFs (Met4 and Met32) that regulate sulfur amino acid pathways (e.g., cysteine, methionine) and is consistent with previous results, under nitrogen limitation, where transcriptional upregulation of methionine and sulfur amino acid pathways was reported (Figure 3G, S3) (Kresnowati et al., 2006). The increase in proteome allocation to vacuole, along with the GO term Fe-S binding discussed above (Figure 3E), was interesting as iron and amino acid homeostasis together with vacuole are considered important for the maintenance of mitochondrial functions that is crucial for the cellular energy budget in yeast (Hughes et al., 2020; Shen, 2020; Weber et al., 2020). In addition, as many vacuolar processes are evolutionary conserved and implicated in diseases such as cancer, diabetes and neurodegeneration further investigation under these conditions could be pertinent for identifying underlying molecular mechanisms in *S. cerevisiae* (Lawrence and Zoncu, 2019; Reggiori and Klionsky, 2013). Overall, our results revealed distinct proteome allocation and synthesis constraints in nutrient-responsive and growth rate-dependent carbon overflow as yeast adapted differently to changes in nutrient environment and growth rate, respectively.

### Energy metabolism trade-off controls proteome abundance and efficiency

As carbon overflow is an integrated readout of cellular metabolism we focused on evaluating the impact of protein abundances (molecules/ pg dcw) on metabolic fluxes in yeast (Figure 4, S1B). First, we evaluated changes in protein abundances constituting the GO terms involved in energy metabolism and protein synthesis, namely glycolysis, the ETC and ribosome (Figure 4A, S2B). We found decrease in the glycolytic protein abundances with the increase in specific glucose uptake rate, except for the critical C:N ratio where total proteome level was maintained similar to the reference but glycolytic abundance increased (Figure 4A, 1C, S1A). A thirty percent increase in the specific glucose uptake rate while maintaining respiration at the critical C:N ratio was achieved differently than 80% increase causing the nutrient responsive-carbon overflow and 385% increase leading to growth rate-dependent carbon overflow indicating diversity in underlying mechanisms causing metabolic overflow (Figure 4A, S1). The glycolytic proteome abundances were likely at their maximum capacity at the critical C:N ratio where increased glycolytic capacity was responsible for increase in the specific glucose uptake rate as compared to the reference (Figure 4A). However, a significant increase in the specific glucose uptake rate, despite a sharp decrease in abundances, under nutrient-responsive and growth rate-dependent carbon overflow was likely achieved by changing other proteome attributes such as the efficiency of glycolytic enzymes (Figure 4A).

**Figure 4:**
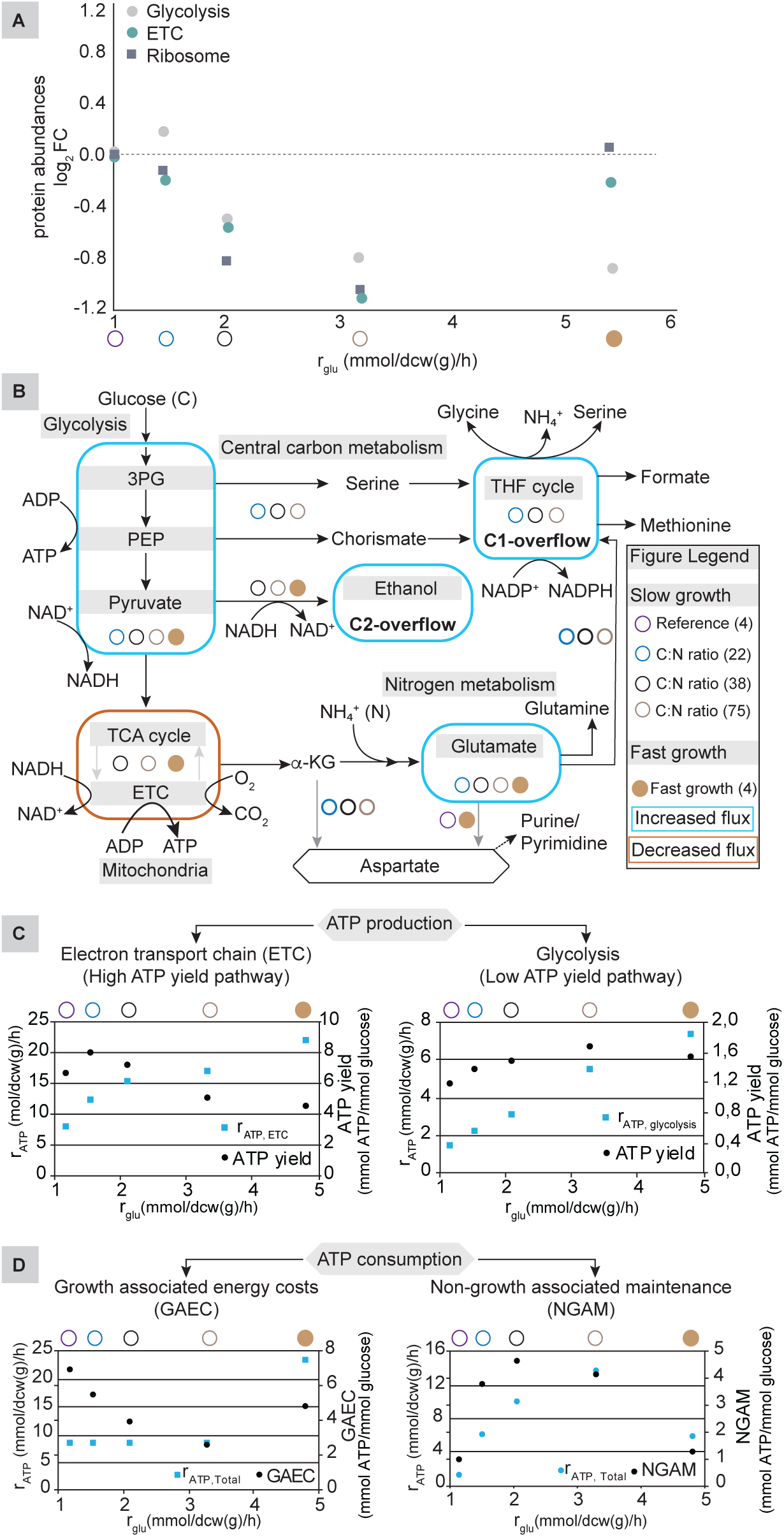
Energy metabolism trade-off controls proteome abundance and efficiency. (A) Relative log_2_ fold changes in protein abundances for the GO terms glycolysis (GO:0006096), ribosome (GO:0005840), and the ETC* compared with the reference on the y-axis are plotted against measured specific glucose uptake rate (mmol/dcw(g)/h) for specified conditions on the x-axis. Each condition is illustrated by an empty or filled circle, description is noted in the figure legend (B). (B) Schematic illustration of changes in the metabolic fluxes in the central carbon and nitrogen metabolism. Figure legend explains design elements of the schematic, where each circle represents an experimental condition and number inside a bracket indicates C:N ratio. (A) Cellular energy efficiency in the high energy (ETC) and low energy (glycolysis) yielding pathways represented by metabolic flux directed at specific ATP production (mmol/dcw(g)/h), shown on primary y-axis, and resulting ATP yield (mmol ATP/ mmol glucose), indicated on secondary y-axis, that are plotted against the specific glycolytic fluxes as determined by the model based on experimental data (measured specific glucose uptake rate) on x-axis. Each condition is illustrated by small empty or filled circle as described (4B). (B) Cellular energy (ATP) expenditure on growth associated energy costs (GAEC) and non-growth associated maintenance (NGAM) - rate (mmol/dcw(g)/h) on primary y-axis and yield (mmol/ mmol glucose) on secondary y-axis. Each condition is illustrated by small empty or filled circle as indicated in the figure legend (B). *Explained in the figure 3D legend See also supplementary S4A, S4B.

Second, to understand the impact of changes in the specific glucose uptake rate and corresponding proteome abundances on metabolism we analyzed the distribution of metabolic fluxes using a *S. cerevisiae* genome-scale metabolic model (GEM) by constraining it with experimentally measured fluxes and maximizing for ATP hydrolysis in metabolism (Figure 4B). This analysis was followed by a flux variability estimation, that allows evaluation of the minimum and maximum range flux for each reaction, using random sampling approach (n=5000) at 95% of the maximal ATP hydrolysis value (Bordel et al., 2010) (Figure 4B, S1B, S4A, S4B). Previously, similar computational approaches have been used to infer changes in efficiency of proteome due to such factors such translation, abundances, metabolites and enzyme catalysis (Bordel et al., 2010; Chen and Nielsen, 2019; Hackett et al., 2016; Tuller et al., 2007). We found that the glycolytic flux increased in synergy with the specific glucose uptake rate, but flux decreased towards the TCA cycle, the ETC and the ATP synthase which together are necessary for aerobic respiration (Figure 4B, S1B, S4A). The increase in the glycolytic flux affected the net contribution of the low ATP yield pathway (glycolysis) and high ATP yield pathway (the ETC) towards cellular energy budget (Figure 4C, S4B). The efficiency of glycolysis measured in terms of its ability to produce ATP increased with the increase in the specific glucose uptake rate despite reduced abundances for the glycolytic proteome at the onset of carbon overflow (Figure 4A-C). The efficiency of the ETC initially increased with the change in the C:N ratio until it reached the critical level where further increase in the C:N ratio reduced the ATP yield in this pathway (Figure 4C). Interestingly, almost similar proteome abundance ratio for ribosome and the ETC was present both at the critical C:N ratio, at the slow growth, and the reference C:N ratio, at the fast growth, but resulted in much lower ATP yield for the latter indicating a reduced ATP contribution from respiration to the cellular energy budget at the fast growth rate (Figure 4C, S4B). This implored us to look at the energy expenditure and we found that the ATP spent on the NGAM was much higher at the critical C:N ratio, at slow growth, compared with the reference C:N ratio at fast growth rate (Figure 4D, S4B). The high NGAM cost might also explain why the critical C:N ratio showed less biomass but similar protein content compared with the reference at the slow rate (Figure 1C, 4D). The efficiency of the ETC was reduced and glycolysis was increased under nitrogen limitation (above the critical C:N ratio) causing nutrient responsive carbon overflow as much of the produced energy was spent on the NGAM instead on the GAEC leading to reduced ribosome abundances and protein content in the biomass (Figure 4A-D, 1B-C, S4B).

Third, we investigated potential pathways used in incepting diverse carbon overflow in yeast (Figure 4B, S1B, S4B). The model predicted different underlying mechanisms causing the nutrient-responsive carbon overflow compared with the growth rate-dependent carbon overflow (Figure 4B, S1B, S4B). It suggested that nitrogen limitation besides activating pathways related to nitrogen metabolism also activated the tetrahydrofolate (THF) cycle (C1 metabolism) affecting folate and methionine biosynthesis together with the precursor metabolite pathways (serine, chorismate) which important as these changes are reported to influence mitochondrial dynamics and play an important role in cancer metabolism (Figure 4B, S1B, S4A) (Gao et al., 2018; Roy et al., 2020). The activation of these pathways led to not just C2 overflow (ethanol) but also C1 overflow (formate) under nitrogen limitation which we validated by measuring formate in culture samples (Figure 4B, S1B, S4A). The likely reason for activation of the THF pathway was redox balance and replenishment of nitrogen (Figure 4B, S1B, S4A). The model also predicted a miniscule carbon overflow of C3 (glycerol, pyruvate) and C5 (2-oxaloactetate) compounds for the critical C:N ratio, a condition which, interestingly, did not show any detectable C2 (ethanol) carbon overflow in our experimental data (Figure 1B, 4B, S1A, S1B). The model predictions were consistent with carbon balance data from bioreactors where about 9% carbon remained missing at the critical C:N ratio and formate measurements allowed carbon balance at higher C:N ratio (Figure 1B, S1B). In the model prediction, the aspartate biosynthesis pathway, that provides backbone for biosynthesis of nucleotides, was also suggested to be C:N ratio dependent where mitochondrial aspartate aminotransferase, Aat1, was preferred at the high C:N ratios (>22) while cytosolic aspartate aminotransferase, Aat2, was utilized at the reference C:N ratios (Figure 4B, S1B, S4A) (Boer et al., 2010). Our results, under nitrogen limitation, obtained using proteome and metabolic modeling response are consistent with previous reports showing, at metabolomics and transcriptome levels, that a drain of adenine pool is accompanied by increase of purine biosynthesis, C1 and sulfur metabolism (Kresnowati et al., 2006).

Finally, our results suggested that the cellular energy budget impinged on composition of biomass (Figure 4, 1C). At the slow growth, even as total proteome abundances decreased due to nitrogen limitation (particularly, ribosome), protein allocation remained similar to the reference (Figure 4, 3A). This resulted in a significant decrease in the total ATP and an energy expenditure trade-off in favor of the NGAM compared with the GAEC causing a nutrient-responsive carbon overflow (Figure 4). At the fast growth, protein allocation was increased for ribosome but decreased for the energy metabolism compared with the reference, though abundances for both the GO terms remained almost similar to the reference (Figure 4, 3A). This led to a reduced efficiency of the ETC due to less available resources for the synthesis of mitochondria and increased efficiency of the glycolysis that was redox balanced through growth rate-dependent carbon overflow (Figure 4, 3A). It was interesting to note that nearly similar protein abundances for ribosomes and the ETC resulted in respiration at the critical C:N ratio condition at slow growth but fermentation at the fast growth condition with the reference C:N ratio (Figure 4A-B). Thus, demonstrating that the fast growth rate-dependent carbon overflow was largely due to the differences in proteome allocation and consequences thereof in metabolism (Figure 4, 3A).

## Discussion

We conducted experiments at a physiological steady state to eliminate confounding effects due to changing cellular environment when cells are cultivated in a batch experiment (Figure 1). This allowed us to infer regulatory effects caused by the changes in nitrogen concentration while keeping a constant supply of glucose and compare these effects at the both slow and fast growth rates in model organism *S. cerevisiae* (Figure 1, S1A). Our systems level analysis showed how energy metabolism controls proteome allocation in growth rate-dependent carbon overflow and proteome abundances in nutrient-responsive carbon overflow in *S. cerevisiae* (Figure 2-4). Our results also showed how trade-off between the low or high ATP yielding pathways determines energy production, and the ATP expenditure towards the GAEC or the NGAM results in either respiration or fermentation mode of metabolism (Figure 4). Finally, we illustrated an important role of balance in cellular energy budget where an imbalance can result in carbon overflow (Figure 5).

**Figure 5:**
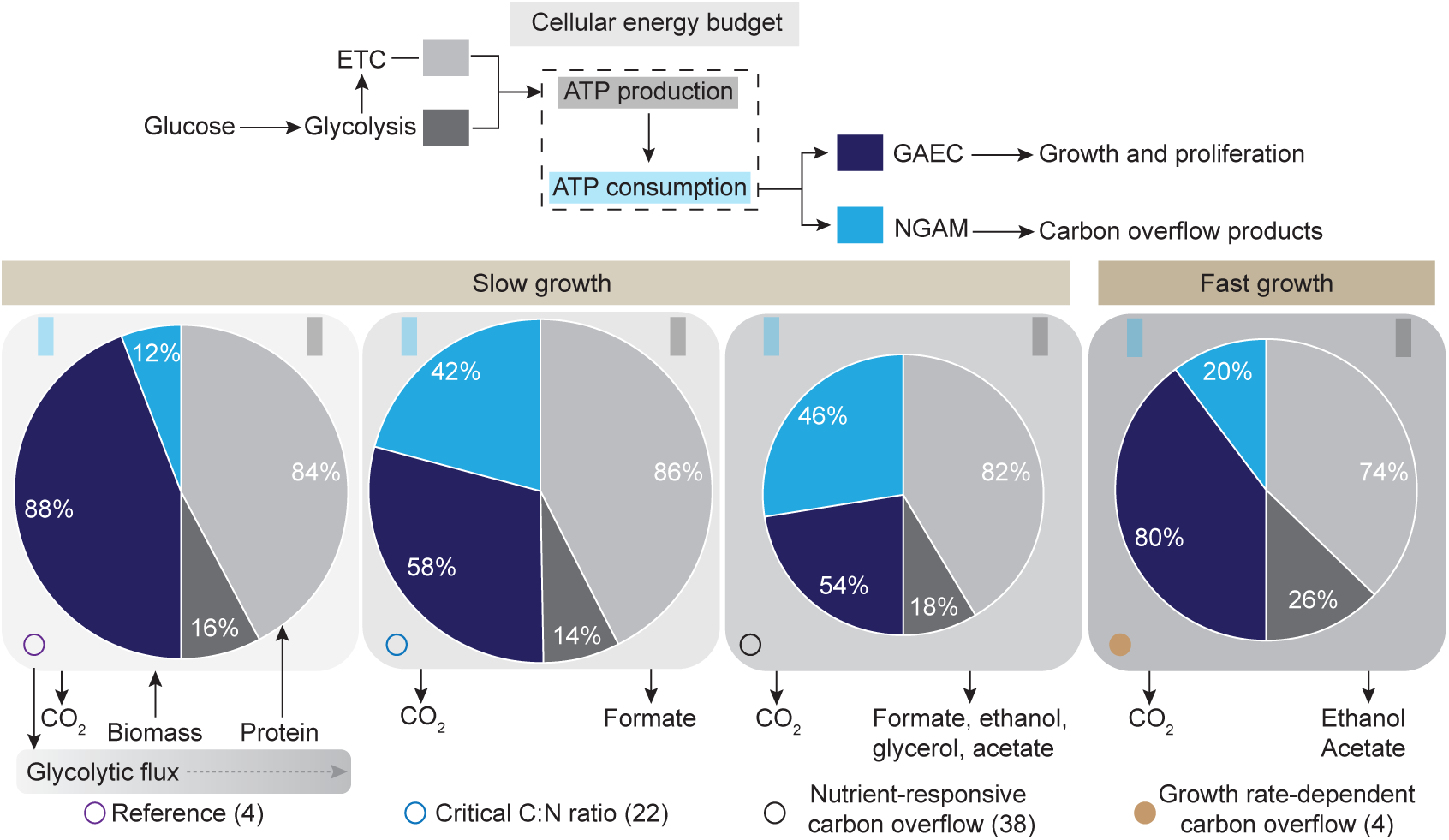
Glycolytic flux impinges on cellular energy budget. Schematic of energy budget showing ATP production and consumption at the cellular level. Top illustration shows input and output of cellular energy budget, and is followed by exhibit of budget distribution at slow and fast growth. Color gradient (light to dark) in the round bounding squares show increase in the glycolytic flux as indicated in the small inset. Impact of energy budget on cellular composition is shown by presence of distinct protein fraction in biomass of each condition and is illustrated by size of pie charts. Numbers on each pie indicate % contribution of each category to cellular energy budget in *S. cerevisiae* (light grey – energy produced in electron transport chain; dark grey – energy produced in glycolysis; light blue – energy spent for non-growth associated maintenance; dark blue – energy spent for biomass formation). Each condition is illustrated by small empty or filled circle as indicated in the figure.

Our results demonstrated that a fully respiratory metabolism represents a cellular energy supply-demand homeostasis at the reference condition (D, 0.1 h^−1^; C:N ratio, 4) where a balance in the catabolic energy supply and anabolic energy demand resulted in maximum production of biomass (Figure 1, S1A). Such homeostasis existed only for C:N ratio 4-10 at the slow growth (D, 0.1 h^−1^) that showed a fully respiratory metabolism (Figure 1, S1A). However, at higher C:N ratio (≥22), at slow growth, and at the reference C:N ratio (4), at the fast growth, the maintenance of homeostasis that allowed for maximum biomass production was not feasible and these conditions resulted in nutrient-responsive and growth rate-dependent carbon overflow, respectively (Figure 1, S1A). These diverse carbon overflows resulted in distinct cellular composition as observed by the percentage change in proteome as a fraction of total biomass (Figure 1). We found that, at the slow growth rate, the maximum ATP efficiency for protein production was not at the reference C:N ratio (4) but was present at the critical C:N ratio (22) (Figure 1B, 4B). It suggests that the chemically defined culture medium used for yeast cultivation, commonly referred to as the Delft medium due to its origin (Verduyn et al., 1992), is rich in nitrogen (5 g/l) but is limited for glucose (10 g/l) at the reference condition and is optimized for maximizing formation of biomass.

We determined the reproducibility of quantitative proteomics data using the PCA and performed the downstream data analysis by using similar statistically relevant filtering approaches for the both, allocation and abundances of proteins as reported in our previous study (Figure 2) (Lahtvee et al., 2017). Here, we found that the differences in the slow and fast growth rates are largely due to differences in cellular resource allocation among the defined GO term categories, namely glycolysis, mitochondrion and ribosome (Figure 3A). At the fast growth, the largest protein allocation shift happens towards ribosome with an increase of 10% compared with the slow growth, but it comes at the cost of allocation towards the energy generating pathways, namely mitochondrion (the ETC) and glycolysis (Figure 3A-C). Therefore, at the slow growth synthesis of mitochondria occurs to the fullest extent possible but at the fast growth rate, a large proteome allocation towards ribosome negatively impacts allocation for mitochondria and thus its contribution towards cellular energy budget (Figure 4). The most striking similarity between the both nutrient-responsive and growth rate-dependent carbon overflow concerns the specific glucose uptake rate that increased almost by 5-fold at the fast growth rate (Figure 4A). At the same time, glycolytic proteins showed a significant decrease in the both allocation and abundances but yielded more ATP due to increase in the specific glucose uptake rate suggesting an increase in efficiency of glycolytic enzymes (Figure 3-4). This suggests that the catalytic efficiency of glycolytic proteins is not determined by allocation or abundance differences but controlled by other factors such as the PTMs as indicated by phosphorylation of glycolytic peptides in our phosphoproteome data (Figure 4, S4A, S2C). This is consistent with the previous report suggesting that glycolysis is mostly regulated at the posttranscriptional levels in *S. cerevisiae* (Daran-Lapujade et al., 2007). As the glycolytic flux increases, more ATP is contributed from the glycolysis compared with the ETC to the cellular energy budget and it is partially because glycolysis can produce more ATP per protein mass as opposed to mitochondria whose synthesis itself is energy intensive process even though overall respiratory chain can produce ATP more efficiently but requires more proteins (Figure 4) (Chen and Nielsen, 2019; Molenaar et al., 2009; Nilsson and Nielsen, 2016). Interestingly, at the slow growth, the critical C:N ratio (22) showed maximum ATP yield by increasing glycolytic protein abundances without significantly reducing protein abundances for the ETC (mitochondrion), but compromising on biomass yield (Figure 1C, 3, 4). Thus, under the condition of slow growth and a relatively low flux the increase in glycolytic protein abundances added spare enzyme capacity to glycolysis (Figure 4). The additional glycolytic enzyme capacity allowed ribosomes to continue the synthesis of mitochondrial proteome almost at the reference level making it near-equilibrium condition for the energy and protein production (Figure 3-4). This observation is consistent with a recent report showing that at relatively slow fluxes multiple steps in glycolysis operate at near equilibrium and reflect spare enzyme capacity (Park et al., 2019). The reduced biomass at the critical ratio was due to higher energy expenditure on the NGAM and less on the GAEC compared with the reference (Figure 4D). At the slow growth and high C:N ratio (>22) the total protein content reduced by nearly 40% compared with the reference (Figure 3). At the C:N ratio (>22) protein allocation for ribosomal proteins remained similar to the critical C:N ratio (22) but reduced by about 2% compared with the reference (4), (Figure 3) indicating that a certain threshold translation capacity was necessary for the maintenance growth rate (0.1 h^−1^). However, at the C:N ratio (>22), ribosome abundances decreased significantly and may explain reduction in total protein content under these conditions as the cellular energy budget mainly relied on glycolysis, reducing mitochondrial synthesis and therefore, the ETC contribution, and spent available energy on the NGAM resulting in nutrient-responsive carbon overflow (Figure 4). This supports the idea that constant growth rate can be maintained by reduced total energy flux by increasing carbon overflow (Slavov et al., 2014). The significant increase in the NGAM at the high C:N ratios suggests activation of cellular homeostasis mechanisms, such as autophagy, to replenish nitrogen by protein turnover as indicated by increased protein allocation for vacuole where previous reports found presence of low intracellular amino acids and high nucleotides in *S. cerevisiae* (Figure 3) (Boer et al., 2010; Marshall et al., 2016). Interestingly, a similar nutrient stress condition also shows the active lysosomal v-ATPase-Ragulator complex, a common activator for AMPK and mTORC1, acting as a switch between catabolism and anabolism in higher eukaryotes (Efeyan et al., 2012; Zhang et al., 2014). From the viewpoint of biotechnology application activation of the C1metabolism and switching of aspartate biosynthesis pathways under nitrogen limitation at slow growth are particularly relevant (Figure 3-4). For example, the THF cycle has been demonstrated for developing bioprocesses in *S. cerevisiae* (Gonzalez de la Cruz et al., 2019). Also, the both C1 metabolism and aspartate biosynthesis are potential anticancer targets as rapidly proliferating mammalian cells can rely upon these metabolites for respiration (Koseki et al., 2018; Meiser et al., 2016; Morscher et al., 2018; Sullivan et al., 2015).

In conclusion, we demonstrated the role of cellular energy budget in respiration, and nutrient-responsive and growth rate-dependent carbon overflow in *S. cerevisiae* (Figure 5). At fast growth, a large resource allocation towards ribosomes constrained resource availability for the synthesis of mitochondrion (respiration) resulting in a growth rate-dependent carbon overflow (Figure 5). At slow growth, total proteome decreased at higher C:N ratios due nitrogen limitation while maintaining a similar cellular resource allocation and growth rate as the reference (Figure 5). The decrease in proteome resulted in a significant reduction in total ATP available for biomass formation which was mostly spent on the NGAM causing a nutrient-responsive carbon overflow (Figure 5). Our results will have practical implications in the both basic science research as well for developing metabolic engineering strategies in biotechnology.

## Materials and Methods

### Strain, media and cultivation conditions

We used yeast *Saccharomyces cerevisiae* CEN.PK113-7D in this research which has been extensively characterized for laboratory research and was also used in our previous studies (Lahtvee et al., 2016; Nijkamp et al., 2012; Zhang et al., 2011). All experiments were performed from the same set of glycerol stocks that were prepared at the beginning of this study. A typical inoculum sample was prepared by reviving an aliquot of glycerol culture stock by passing through two pre-culture stages in a chemical defined medium before being collected at the exponential phase to be used in an experiment. The chemically defined medium, commonly referred to as the Delft medium in laboratories due to origin (Verduyn et al., 1992), used contained per liter glucose (20 g in pre-culture and batch bioreactors; 10 g in feed medium of chemostat experiments), ammonium sulfate (5 g), potassium dihydrogen phosphate (3 g) and magnesium sulfate heptahydrate (0.5 g). It was supplemented with 1 ml each of trace metals and vitamins stock solutions similar to our previous studies (Lahtvee et al., 2016; Zhang et al., 2011). In bioreactor experiments 50 μl of Antifoam 204 (Sigma-Aldrich, USA) per liter was added to the culture medium. All experiments were conducted in triplicate using 1 l bioreactors (Applikon BIOTECHNOLOGY) with a 0.5 l working volume. The bioreactors controls were used to maintain a constant volume, pH (at 5.5 by use of 2 M KOH), and fully aerobic conditions (1 volume per volume per minute, vvm) throughout the experiments. Dilution rates (D) were used to control growth rates in chemostat experiments and data were collected after at least three residence times (1/D) passed at steady state which was monitored for stability by online sensors for off-gas (CO_2_ and residual O_2_), dissolved oxygen (DO) culture medium, and biomass (also cross-checked offline). Chemostat experiments were performed on an average of six residence times and samples for analysis were collected at a steady state only after lapse of at least three residence times. A steady state was determined by constancy of gaseous exchanges online and optical density/biomass measurements offline. In all chemostat experiments glucose concentration (carbon) was aimed to be constant while ammonium sulfate (nitrogen) concentration was varied to obtain a range of C:N ratios. Biological triplicate data were collected for dry cell weight (biomass), metabolites, quantitative proteome and phosphoproteome as described in our previous publications (Lahtvee et al., 2017; Zhang et al., 2011).

### Metabolites measurements

Culture broth samples from the steady state experiments and feed bottles were collected in sterile Eppendorf tubes (2 ml) for quantification of extracellular metabolites. The collected samples were centrifuged at 14,800 rpm at 4^0^C for 10 min to remove biomass and stored at -20^0^C until analysis using a high-performance liquid chromatography (HPLC). Extracellular metabolites were quantified using an Aminex HPX-87H chromatography column (Bio-Rad) with the recommended settings for elution of sugars and organic acids (temperature 45^0^C; flow rate 0.6 ml/min using mobile phase of 5 mM sulfuric acid) in a refractive index and UV detector containing HPLC instrument (Prominence-I, LC-2030 C Plus, SHIMADZU).

### Total yeast proteome: extraction and quantification

All biomass samples were normalized to equal concentrations (1 g/l) and 600 μg of biomass per sample was resuspended in a commercially available protein extraction solution (Y-PER^TM^ Yeast, Thermofisher). This biomass suspension in a vial (2 ml, Eppendorf) was thoroughly mixed with pipette and all vials were incubated at 30^0^C for 45 min under moderate agitation conditions. After incubation biomass suspension from each vial was transferred to a glass-beads containing cells lysing tube (Precellys). The cells lysing tubes were repeatedly agitated for 10 cycles, with an interval of 5 min after each cycle (4 m/s for 20 s), in a FastPrep-24 device. These agitated tubes were centrifuged at 14,800 rpm at 4^0^C for 10 min and supernatant was carefully removed from each tube. A fraction of biomass remained leftover amid glass beads in the lysing tubes and an equal volume of Y-PER reagent was added to repeat (without 45 min incubation period) disruption and extraction using a FastPrep-24 device. This step was repeated until total proteome was extracted from yeast biomass. For each step proteome was quantified using a commercially available assay kit (Micro BCA^TM^ Protein Assay Kit, Thermofisher) and samples were diluted to be in the linear range of BSA protein standard range (0.5 to 20 μg/ml). Proteome assays were performed in triplicate to ensure reproducibility of data and cumulative proteome obtained from each step reflects total proteome of yeast.

### Mass-spectrometric quantification of proteome and phosphoproteome

Quantification of absolute proteome and identification of phosphoproteome in samples was done similar to our previous study using a Nano-LC/MS-MS analysis in combination with extensively used MaxQuant 1.4.08 software package (Cox and Mann, 2008; Lahtvee et al., 2017). Briefly, modifications to our previous studies on both quantification of absolute proteome and phosphoproteome are described here (Lahtvee et al., 2017; Zhang et al., 2011). Sample biomass pellets were lysed in glass beads containing Eppendorf tubes at pH 8.0 buffer (6 M guanidine HCl, 100 mM Tris-HCl, 20 mM dithiothreitol) and homogenized using FastPrep24 (MB Biomedicals) cells disruptor with two cycles (4 m/s for 30s). Supernatant was removed by centrifuge (17000 g, 10 min, 4^0^C), precipitated overnight with 10% trichloroacetic acid (TCA) at 4^0^C, and assayed for protein concentration as described above in the total yeast proteome section. Both absolute protein quantification and phosphoproteome sample preparations were similar to previous descriptions except phosphoproteome samples were not mixed with heavy standards and were digested with trypsin instead of Lys-C enzyme (Humphrey et al., 2015; Lahtvee et al., 2017). For phosphoproteome enrichment 500 μg of sample protein was used and reconstituted in 0.5% trifluoroacetic acid (TFA) similar to samples for quantification of absolute proteome. For phosphoproteome total enriched sample and for absolute quantification 2 μg of protein sample were used in Nano-LC/MS-MS analysis (Lahtvee et al., 2017). Peptides were separated at 200 nL/min for absolute proteome and 250 nL/min for phosphoproteome with a 5-40% B 240 min gradient for spiked time point and 480 min gradient for heavy standard samples. For phosphopeptides a 90 min separating gradient was used in 5-15% for 60 min and 15-30% for 30 min steps. Normalized collision energies of 26 for normal peptides and 27 for phosphopeptides were used in higher-energy collisional fragmentation process. For absolute proteome analysis the ion target and injection times for the MS were 3× 10^6^ (50 ms) while for the MS/MS were 3× 10^6^ (50 ms). The same parameters for phosphopeptides were at 1× 10^6^ (60 ms) and 2× 10^4^ (60 ms), respectively. Dynamic exclusion time was limited to 110, 70 and 45s for heavy standard samples, spiked time point and phosphopeptides, respectively. Only charge states +2 and +6 were targeted to MS/MS and additionally a fixed first mass of 95 m/z was set for phosphopeptides. All heavy standards were analyzed as technical triplicates while single replicate analysis was performed for biological triplicate samples. For phospho-analysis serine/threonine phosphorylation was used as additional variable modification in addition to previously described variable modifications for absolute proteome quantification (Lahtvee et al., 2017). *Saccharomyces cerevisiae* reference proteome database (version July 2016) was accessed at the UniProt (www.uniprot.org) and was searched using the LysC/P (absolute proteome) and the trypsin/P (phosphoproteome) digestion rules. Raw data quantification was carried out by dividing protein intensities from heavy standard with the number of theoretically observable peptides, log-transformed and plotted against log-transformed UPS2 mix (48 human proteins) with known protein abundances. This regression was then used to derive all other protein absolute quantities using their iBAQ intensities. Normalized H/L ratios (by shifting median peptide log H/L ratio to zero) were used for all downstream quantitative analyses (Cox and Mann, 2008). For biomass normalized and protein normalized absolute quantities, total protein amount in dcw (g) was either considered or not, respectively. This resulted in quantities of molecules of individual protein per pg dcw in case of biomass normalized data, and molecules of individual protein per pg of total protein in sample in case of protein normalized data. The mass spectrometry proteomics data have been deposited to the ProteomeXchange Consortium via the PRIDE partner repository with the dataset identifier PXD016854 (Perez-Riverol et al., 2019).

### Data analysis pipeline for absolute proteome and phosphoproteome

Differential expression analysis for absolute proteins and phosphoproteome was carried out in R, statistical differences were calculated using Student’s t-test and false discovery rate (FDR) according to Benjamini–Hochberg procedure. Gene set analysis was performed with the biomass and protein normalized proteomics data using Piano, with the mean of the gene-level statistics, ignoring gene-sets smaller than 5 and larger than 500 genes, sampling 5,000 times and corrected for multiple testing using FDR (Varemo et al., 2013). Same platform was used for TF analysis, with TF-gene relationships acquired from the yeastract (http://www.yeastract.com). Principal Component Analysis was carried out in ClustVis (Metsalu and Vilo, 2015). A response-decomposition framework developed for further analysis of this proteomics data is described in the Supplemental M.

### Flux balance analysis (FBA)

Intracellular distribution of metabolic fluxes was investigated with the *S. cerevisiae* genome-scale metabolic model yeast-GEM version 8.3.4 (Sánchez et al., 2019). Calculations were performed with Cobra Toolbox (Schellenberger et al., 2011) on MATLAB (The MathWorks Inc., Natick, MA, USA) using Gurobi solver (Gurobi Optimization Inc., Houston, TX, USA). First, ATP hydrolysis, representing non-growth associated maintenance energy, was maximized to calculate the unique pattern of intracellular fluxes. To determine the variability of fluxes, random sampling algorithm in RAVEN was used with 5000 samplings at the 95% previously determined ATP drain value (Bordel et al., 2010). This resulted in an average flux with the standard deviation, representing the flux variability.

## Supporting information

S1A_Physiology

S1B_Fluxes

S2A_Prot_allocation

S2B_Prot_abundance

S2C_Phosphoproteome

S2D_Enrichment

S3_GSA

S4A_Metabolic Pathways

S4B_Cellular energy budget

Supplemental M - Proteomics Data Analysis Method

## Acknowledgment

We would like to thank Dr. Kaspar Valgepea, TUIT, Estonia and Dr. Nicolaas A Buijs, Chr. Hansen A/S, Denmark, for feedback on the manuscript and TUIT Synbio lab members for valuable support during the project.

## Supplementary Captions

**S1A:** Experimental data generated using a chemical defined culture medium (described in the manuscript). Table metadata is provided in the first two columns of this sheet.

**S1B:** Flux calculations with the yeast S*accharomyces cerevisiae* genome-scale model version 8.3.4. Flux balance analysis was carried out by constraining measured uptake and production fluxes (including specific growth rate), while optimizing for ATP synthesis (r_0226), followed by constraining ATP synthesis at 95% of the maximal value and running random sampling algorithm (n, 5000) for determination of flux variability. Average flux values from random sampling together with its standard deviation are presented. ID - reaction identification number in the model; Name - reaction name; EC - number - EC number corresponding to the reaction; Gene Association - reaction associated genes; REF - reference chemostat experiment (D, 0.1 h^−1^, C:N molar ratio of 4); CN22 - chemostat at D, 0.1 h^−1^, C:N molar ratio of 22; CN38 - chemostat at D, 0.1 h^−1^, C:N molar ratio of 38; CN75 - chemostat at D, 0.1 0.1 h^−1^, C:N molar ratio of 75; Fast growth (FG) - experiments at elevated dilution rates (D, 0.26 or 0.32 h^−1^, C:N molar ratio of 4).

**S2A:** Amount of proteome in *Saccharomyces cerevisiae* chemostat experiments reported in molecules/pg-protein (used in results/discussion on “protein allocation”). REF - reference chemostat experiment (D, 0.1 h^−1^, C:N molar ratio of 4); CN22 - chemostat at D, 0.1 h^−1^, C:N molar ratio of 22; CN38 - chemostat at D, 0.1 h^−1^, C/N molar ratio of 38; CN75 - chemostat at D, 0.1 h^−1^, C:N molar ratio of 75; Fast growth (FG) -experiments at elevated dilution rates (D, 0.26 or 0.32 h^−1^, C:N molar ratio of 4). Adjusted p-values were calculated using the Benjamini-Hochberg procedure.

**S2B.** Protein abundances in *Saccharomyces cerevisiae* chemostat experiments reported in molecules/pg- dcw (used in results/discussion on “protein abundances”). REF - reference chemostat experiment (D, 0.1 h^−1^, C:N molar ratio of 4); CN22 - chemostat at D, 0.1 h^−1^, C:N molar ratio of 22; CN38 - chemostat at D, 0.1 h^−1^, C:N molar ratio of 38; CN75 - chemostat at D, 0.1 h^−1^, C:N molar ratio of 75; Fast growth (FG) -experiments at elevated dilution rates (D, 0.26 or 0.32 h^−1^, C:N molar ratio of 4).

**S2C:** Phosphoproteome in *Saccharomyces cerevisiae* chemostat experiments. REF - reference chemostat experiment (D, 0.1 h^−1^, C:N molar ratio of 4); CN22 - chemostat at D, 0.1 h^−1^, C:N molar ratio of 22; CN38 - chemostat at D, 0.1 h^−1^, C:N molar ratio of 38; CN75 - chemostat at D, 0.1 h^−1^, C:N molar ratio of 75; Fast growth (FG) -experiments at elevated dilution rates (D, 0.26 or 0.32 h^−1^, C:N molar ratio of 4).

**S2D:** Gene-set analysis using total quantitative proteome and phosphoproteome data to determine representative functional groups in each category.

**S3:** Gene-set analysis using absolute quantitative proteome data. (A) biomass normalized, (B) proteome normalized, (C) summary of significant GO terms and (D) summary of significant transcription factors (TFs)

**S4A:** An overview of central metabolism and pathways responding to changes in C:N ratio vs fast growth driven carbon overflow based on metabolic flux (mmol/g dcw/h) analysis with overlay of absolute proteome abundances and phosphoproteome data. Absolute proteome abundances are indicated for the reference by numbers while relative log_2_ fold changes (biomass normalized) are represented as heatmaps for C:N ratio (22), C:N ratio (38) and fast growth (4). The figure legend explains design elements of illustration. The directionality of metabolic pathways is based on the metabolic flux balance analysis. In central carbon metabolism, at slow growth (D, 0.1 h^−1^), critical C:N ratio (22) showed an increase or similarity to reference abundances while a sharp decrease for most abundances was noticed on further limitation of nitrogen at C:N ratio (38) that was similar to fast growth (C:N ratio, 4) driven carbon overflow. At critical C:N ratio (22) only C1 carbon (formate) overflow was observed but not that of C2 carbon i.e., ethanol. Overflow of C1 carbon, through the THF cycle, was observed in nitrogen limiting conditions but was found absent at the reference C:N ratio (4). Glycolytic proteome showed reduced abundances at the onset of overflow metabolism, but in response to nitrogen assimilation pathway abundances showed an increase where later was also observed for critical C:N ratio (4). Based on metabolic flux analysis, a reference vs high C:N ratio distinction was observed for the biosynthesis of aspartate. Protein abundances for enzymes involved in serine biosynthesis, mitochondrial glycine decarboxylation and methionine biosynthesis increased at critical C:N ratio (22). Abbreviations: G-6-P, glucose 6 phosphate; F-6-P, fructose 6 phosphate; F-1,6-P, fructose 1,6 bisphosphate; DHAP, dihydroxyacetone phosphate; G-3-P, glyceraldehyde 3 phosphate; 3PGP, 3 phosphoglyceroyl phosphate;3PG, 3 phosphoglycerate; 2PG, 2 phosphoglycerate; PEP, phosphoenolpyruvate; FAs, fatty acids; OAA, oxaloacetic acid; CIT, citric acid; ICIT, iso-citric acid; a-KG, alpha-ketoglutaric acid; SUC- CoA, succinyl CoA; SUC, succinic acid; FUM, fumaric acid; MAL, malic acid; 7,8-DHF, a 7,8 dihydrofolate; 5,10-mTHF, a 5,10 methylene tetrahydrofolate; 5-mTHF, 5 methyl tetrahydrofolate; 10- fTHF, a 10 formyl tetrahydrofolate; THF, a tetrahydrofolate; NA, Protein abundance not measured. Reaction number corresponds to corresponding metabolic reactions in the supplementary S1B. Reaction number (metabolic reactions, S1B): 1 (r_0534), 2 (r_0467), 3 (r_0886, r_0449), 4 (r_0450), 5 (r_1054), 6 (r_0486), 7 (r_0892), 8 (r_0893), 9 (r_0366), 10 (r_0962), 11 (r_0959, r_0960), 12 (r_0173), 13 (r_0112), 14 (r_2140, 2141), 15 (r_2115), 16 (r_0891), 17 (r_0961), 18 (r_0958), 19 (r_0719), 20 (r_0300), 21 (r_2305, r_0542, r_4262), 22 (r_0658), 23 (r_0831, r_0832, r_0505), 24 (r_1022, r_0688), 25 (r_1021), 26 (r_0452), 27 (r_0713), 28 (r_0217), 29 (r_0216), 30 (r_0471), 31 (r_0476), 32 (r_0350, r_0347), 33 (r_0997), 34 (r_0066), 35 (r_1045), 36 (r_0725, r_0732, r_0446), 37 (r_0501, r_505), 38 (r_0503), 39 (r_0502), 40 (r_0080).

**S4B:** Cellular energy budget data underlying Figure 4C and 4D. Both metadata and data are described in the table.

**Supplemental M:** A response-decomposition framework for proteomics data analysis

## References

Basan, M., Hui, S., Okano, H., Zhang, Z., Shen, Y., Williamson, J.R., and Hwa, T. (2015). Overflow metabolism in Escherichia coli results from efficient proteome allocation. Nature 528, 99–104.

Boer, V.M., Crutchfield, C.A., Bradley, P.H., Botstein, D., and Rabinowitz, J.D. (2010). Growth-limiting intracellular metabolites in yeast growing under diverse nutrient limitations. Molecular Biology of the Cell 21, 198–211.

Bordel, S., Agren, R., and Nielsen, J. (2010). Sampling the solution space in genome-scale metabolic networks reveals transcriptional regulation in key enzymes. PLoS Comput Biol 6, e1000859.

Broach, J.R. (2012). Nutritional control of growth and development in yeast. Genetics 192, 73–105.

Buijs, N.A., Zhou, Y.J., Siewers, V., and Nielsen, J. (2015). Long-chain alkane production by the yeast Saccharomyces cerevisiae. Biotechnol Bioeng 112, 1275–1279.

Canelas, A.B., Harrison, N., Fazio, A., Zhang, J., Pitkanen, J.P., van den Brink, J., Bakker, B.M., Bogner, L., Bouwman, J., Castrillo, J.I., et al. (2010). Integrated multilaboratory systems biology reveals differences in protein metabolism between two reference yeast strains. Nat Commun 1, 145.

Chen, Y., and Nielsen, J. (2019). Energy metabolism controls phenotypes by protein efficiency and allocation. Proceedings of the National Academy of Sciences of the United States of America 116, 17592–17597.

Cox, J., and Mann, M. (2008). MaxQuant enables high peptide identification rates, individualized p.p.b.-range mass accuracies and proteome-wide protein quantification. Nature Biotechnology 26, 1367–1372.

Crabtree, H.G. (1928). The carbohydrate metabolism of certain pathological overgrowths. Biochem J 22, 1289–1298.

Daran-Lapujade, P., Rossell, S., van Gulik, W.M., Luttik, M.A.H., de Groot, M.J.L., Slijper, M., Heck, A.J.R., Daran, J.M., de Winde, J.H., Westerhoff, H.V., et al. (2007). The fluxes through glycolytic enzymes in Saccharomyces cerevisiae are predominantly regulated at posttranscriptional levels. Proceedings of the National Academy of Sciences of the United States of America 104, 15753–15758.

De Deken, R.H. (1966). The Crabtree effect: a regulatory system in yeast. J Gen Microbiol 44, 149–156.

Efeyan, A., Zoncu, R., and Sabatini, D.M. (2012). Amino acids and mTORC1: from lysosomes to disease. Trends Mol Med 18, 524–533.

Gao, X., Lee, K., Reid, M.A., Sanderson, S.M., Qiu, C.P., Li, S.Q., Liu, J., and Locasale, J.W. (2018). Serine Availability Influences Mitochondrial Dynamics and Function through Lipid Metabolism. Cell Reports 22, 3507–3520.

Gonzalez de la Cruz, J., Machens, F., Messerschmidt, K., and Bar-Even, A. (2019). Core Catalysis of the Reductive Glycine Pathway Demonstrated in Yeast. ACS Synth Biol 8, 911–917.

Hackett, S.R., Zanotelli, V.R., Xu, W., Goya, J., Park, J.O., Perlman, D.H., Gibney, P.A., Botstein, D., Storey, J.D., and Rabinowitz, J.D. (2016). Systems-level analysis of mechanisms regulating yeast metabolic flux. Science 354, 6311.

Hughes, C.E., Coody, T.K., Jeong, M.Y., Berg, J.A., Winge, D.R., and Hughes, A.L. (2020). Cysteine Toxicity Drives Age-Related Mitochondrial Decline by Altering Iron Homeostasis. Cell 180, 296–310 e218.

Humphrey, S.J., Azimifar, S.B., and Mann, M. (2015). High-throughput phosphoproteomics reveals in vivo insulin signaling dynamics. Nat Biotechnol 33, 990–995.

Koseki, J., Konno, M., Asai, A., Colvin, H., Kawamoto, K., Nishida, N., Sakai, D., Kudo, T., Satoh, T., Doki, Y., et al. (2018). Enzymes of the one-carbon folate metabolism as anticancer targets predicted by survival rate analysis. Scientific reports 8, 303.

Kresnowati, M.T., van Winden, W.A., Almering, M.J., ten Pierick, A., Ras, C., Knijnenburg, T.A., Daran-Lapujade, P., Pronk, J.T., Heijnen, J.J., and Daran, J.M. (2006). When transcriptome meets metabolome: fast cellular responses of yeast to sudden relief of glucose limitation. Molecular systems biology 2, 49.

Kumar, R., Lahtvee, P.-J., and Nielsen, J. (2014). Systems Biology: Developments and Applications. In Molecular Mechanisms in Yeast Carbon Metabolism, J. Piškur, and C. Compagno, eds. (Berlin, Heidelberg: Springer Berlin Heidelberg), pp. 83–96.

Kumar, R., and Shimizu, K. (2010). Metabolic regulation of Escherichia coli and its gdhA, glnL, gltB, D mutants under different carbon and nitrogen limitations in the continuous culture. Microb Cell Fact 9, 8.

Kumar, R., and Shimizu, K. (2011). Transcriptional regulation of main metabolic pathways of cyoA, cydB, fnr, and fur gene knockout Escherichia coli in C-limited and N-limited aerobic continuous cultures. Microb Cell Fact 10, 3.

Lahtvee, P.J., Kumar, R., Hallstrom, B.M., and Nielsen, J. (2016). Adaptation to different types of stress converge on mitochondrial metabolism. Molecular biology of the cell 27, 2505–2514.

Lahtvee, P.J., Sanchez, B.J., Smialowska, A., Kasvandik, S., Elsemman, I.E., Gatto, F., and Nielsen, J. (2017). Absolute Quantification of Protein and mRNA Abundances Demonstrate Variability in Gene-Specific Translation Efficiency in Yeast. Cell systems 4, 495–504 e495.

Larsson, C., von Stockar, U., Marison, I., and Gustafsson, L. (1993). Growth and metabolism of Saccharomyces cerevisiae in chemostat cultures under carbon-, nitrogen-, or carbon- and nitrogen-limiting conditions. J Bacteriol 175, 4809–4816.

Lawrence, R.E., and Zoncu, R. (2019). The lysosome as a cellular centre for signalling, metabolism and quality control. Nat Cell Biol 21, 133–142.

Marshall, R.S., McLoughlin, F., and Vierstra, R.D. (2016). Autophagic Turnover of Inactive 26S Proteasomes in Yeast Is Directed by the Ubiquitin Receptor Cue5 and the Hsp42 Chaperone. Cell Reports 16, 1717–1732.

Martin-Perez, M., and Villen, J. (2017). Determinants and Regulation of Protein Turnover in Yeast. Cell systems 5, 283–294 e285.

Meiser, J., Tumanov, S., Maddocks, O., Labuschagne, C.F., Athineos, D., Van Den Broek, N., Mackay, G.M., Gottlieb, E., Blyth, K., Vousden, K., et al. (2016). Serine one-carbon catabolism with formate overflow. Sci Adv 2–10, e1601273.

Metsalu, T., and Vilo, J. (2015). ClustVis: a web tool for visualizing clustering of multivariate data using Principal Component Analysis and heatmap. Nucleic Acids Res 43–W1, W566-570.

Metzl-Raz, E., Kafri, M., Yaakov, G., Soifer, I., Gurvich, Y., and Barkai, N. (2017). Principles of cellular resource allocation revealed by condition-dependent proteome profiling. Elife 6:e28034.

Molenaar, D., van Berlo, R., de Ridder, D., and Teusink, B. (2009). Shifts in growth strategies reflect tradeoffs in cellular economics. Molecular systems biology 5, 323.

Morscher, R.J., Ducker, G.S., Li, S.H., Mayer, J.A., Gitai, Z., Sperl, W., and Rabinowitz, J.D. (2018). Mitochondrial translation requires folate-dependent tRNA methylation. Nature 554, 128–132.

Nijkamp, J.F., van den Broek, M., Datema, E., de Kok, S., Bosman, L., Luttik, M.A., Daran-Lapujade, P., Vongsangnak, W., Nielsen, J., Heijne, W.H., et al. (2012). De novo sequencing, assembly and analysis of the genome of the laboratory strain Saccharomyces cerevisiae CEN.PK113–7D, a model for modern industrial biotechnology. Microb Cell Fact 11, 36.

Nilsson, A., and Nielsen, J. (2016). Metabolic Trade-offs in Yeast are Caused by F1F0-ATP synthase. Scientific reports 6, 22264.

Oliveira, A.P., and Sauer, U. (2012). The importance of post-translational modifications in regulating Saccharomyces cerevisiae metabolism. FEMS Yeast Res 12, 104–117.

Park, J.O., Tanner, L.B., Wei, M.H., Khana, D.B., Jacobson, T.B., Zhang, Z., Rubin, S.A., Li, S.H., Higgins, M.B., Stevenson, D.M., et al. (2019). Near-equilibrium glycolysis supports metabolic homeostasis and energy yield. Nat Chem Biol 15, 1001–1008.

Peebo, K., Valgepea, K., Maser, A., Nahku, R., Adamberg, K., and Vilu, R. (2015). Proteome reallocation in Escherichia coli with increasing specific growth rate. Mol Biosyst 11, 1184–1193.

Perez-Riverol, Y., Csordas, A., Bai, J., Bernal-Llinares, M., Hewapathirana, S., Kundu, D.J., Inuganti, A., Griss, J., Mayer, G., Eisenacher, M., et al. (2019). The PRIDE database and related tools and resources in 2019: improving support for quantification data. Nucleic Acids Res 47, D442–D450.

Reggiori, F., and Klionsky, D.J. (2013). Autophagic processes in yeast: mechanism, machinery and regulation. Genetics 194, 341–361.

Roy, D.G., Chen, J., Mamane, V., Ma, E.H., Muhire, B.M., Sheldon, R.D., Shorstova, T., Koning, R., Johnson, R.M., Esaulova, E., et al. (2020). Methionine Metabolism Shapes T Helper Cell Responses through Regulation of Epigenetic Reprogramming. Cell Metab 31, 250–266 e259.

Sánchez, B., and, F.L., Lu, H., Kerkhoven, E., and Nielsen, J. (2019). SysBioChalmers/yeast-GEM: yeast 8.3.4. Zenodo v8.3.4.

Schellenberger, J., Que, R., Fleming, R.M., Thiele, I., Orth, J.D., Feist, A.M., Zielinski, D.C., Bordbar, A., Lewis, N.E., Rahmanian, S., et al. (2011). Quantitative prediction of cellular metabolism with constraint-based models: the COBRA Toolbox v2.0. Nat Protoc 6, 1290–1307.

Shen, H. (2020). An IRON-clad Connection between Aging Organelles. Cell 180, 214–216.

Shimizu, K., and Matsuoka, Y. (2019). Regulation of glycolytic flux and overflow metabolism depending on the source of energy generation for energy demand. Biotechnol Adv 37, 284–305.

Slavov, N., Budnik, B.A., Schwab, D., Airoldi, E.M., and van Oudenaarden, A. (2014). Constant growth rate can be supported by decreasing energy flux and increasing aerobic glycolysis. Cell Rep 7, 705–714.

Sullivan, L.B., Gui, D.Y., Hosios, A.M., Bush, L.N., Freinkman, E., and Vander Heiden, M.G. (2015). Supporting Aspartate Biosynthesis Is an Essential Function of Respiration in Proliferating Cells. Cell 162, 552–563.

Torrence, M.E., and Manning, B.D. (2018). Nutrient Sensing in Cancer. Annual Review of Cancer Biology, Vol 2 *2*, 251–269.

Tuller, T., Kupiec, M., and Ruppin, E. (2007). Determinants of protein abundance and translation efficiency in S-cerevisiae. Plos Computational Biology 3, 2510–2519.

Vander Heiden, M.G., Cantley, L.C., and Thompson, C.B. (2009). Understanding the Warburg effect: the metabolic requirements of cell proliferation. Science 324, 1029–1033.

Varemo, L., Nielsen, J., and Nookaew, I. (2013). Enriching the gene set analysis of genome-wide data by incorporating directionality of gene expression and combining statistical hypotheses and methods. Nucleic Acids Res 41, 4378–4391.

Verduyn, C., Postma, E., Scheffers, W.A., and Van Dijken, J.P. (1992). Effect of benzoic acid on metabolic fluxes in yeasts: a continuous-culture study on the regulation of respiration and alcoholic fermentation. Yeast 8, 501–517.

Vlastaridis, P., Papakyriakou, A., Chaliotis, A., Stratikos, E., Oliver, S.G., and Amoutzias, G.D. (2017). The Pivotal Role of Protein Phosphorylation in the Control of Yeast Central Metabolism. G3 (Bethesda, Md) 7, 1239–1249.

Warburg, O. (1956). On the Origin of Cancer Cells. Science 123, 3191.

Weber, R.A., Yen, F.S., Nicholson, S.P.V., Alwaseem, H., Bayraktar, E.C., Alam, M., Timson, R.C., La, K., Abu-Remaileh, M., Molina, H., et al. (2020). Maintaining Iron Homeostasis Is the Key Role of Lysosomal Acidity for Cell Proliferation. Mol Cell 77, 645–655 e647.

Yu, T., Zhou, Y.J., Huang, M., Liu, Q., Pereira, R., David, F., and Nielsen, J. (2018). Reprogramming Yeast Metabolism from Alcoholic Fermentation to Lipogenesis. Cell 174, 1549–1558 e1514.

Zhang, C.S., Jiang, B., Li, M., Zhu, M., Peng, Y., Zhang, Y.L., Wu, Y.Q., Li, T.Y., Liang, Y., Lu, Z., et al. (2014). The lysosomal v-ATPase-Ragulator complex is a common activator for AMPK and mTORC1, acting as a switch between catabolism and anabolism. Cell Metab 20, 526–540.

Zhang, J., Vaga, S., Chumnanpuen, P., Kumar, R., Vemuri, G.N., Aebersold, R., and Nielsen, J. (2011). Mapping the interaction of Snf1 with TORC1 in Saccharomyces cerevisiae. Molecular systems biology 7, 545.

