## Supplementary figures and images for "Proteome overabundance enables respiration but limitation onsets carbon overflow"

### S2D_Enrichment

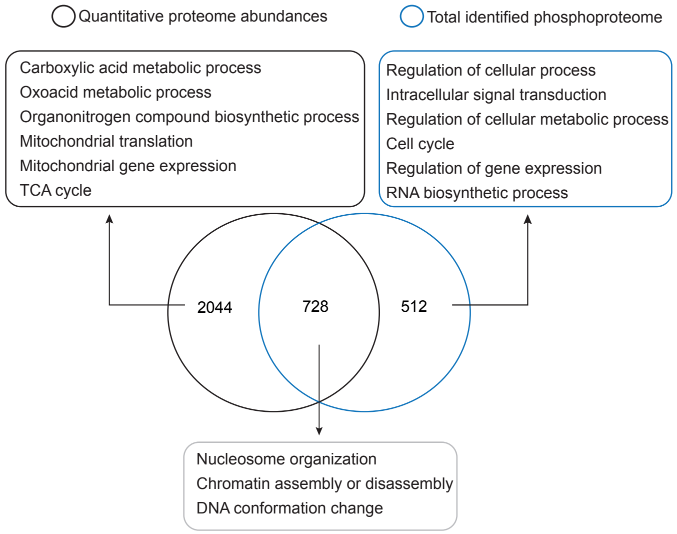


S2D: Gene-set enrichment analysis using quantitative proteome and phosphoproteome data

### S3_GSA

S3. Gene-set analysis using quantitative proteomics data

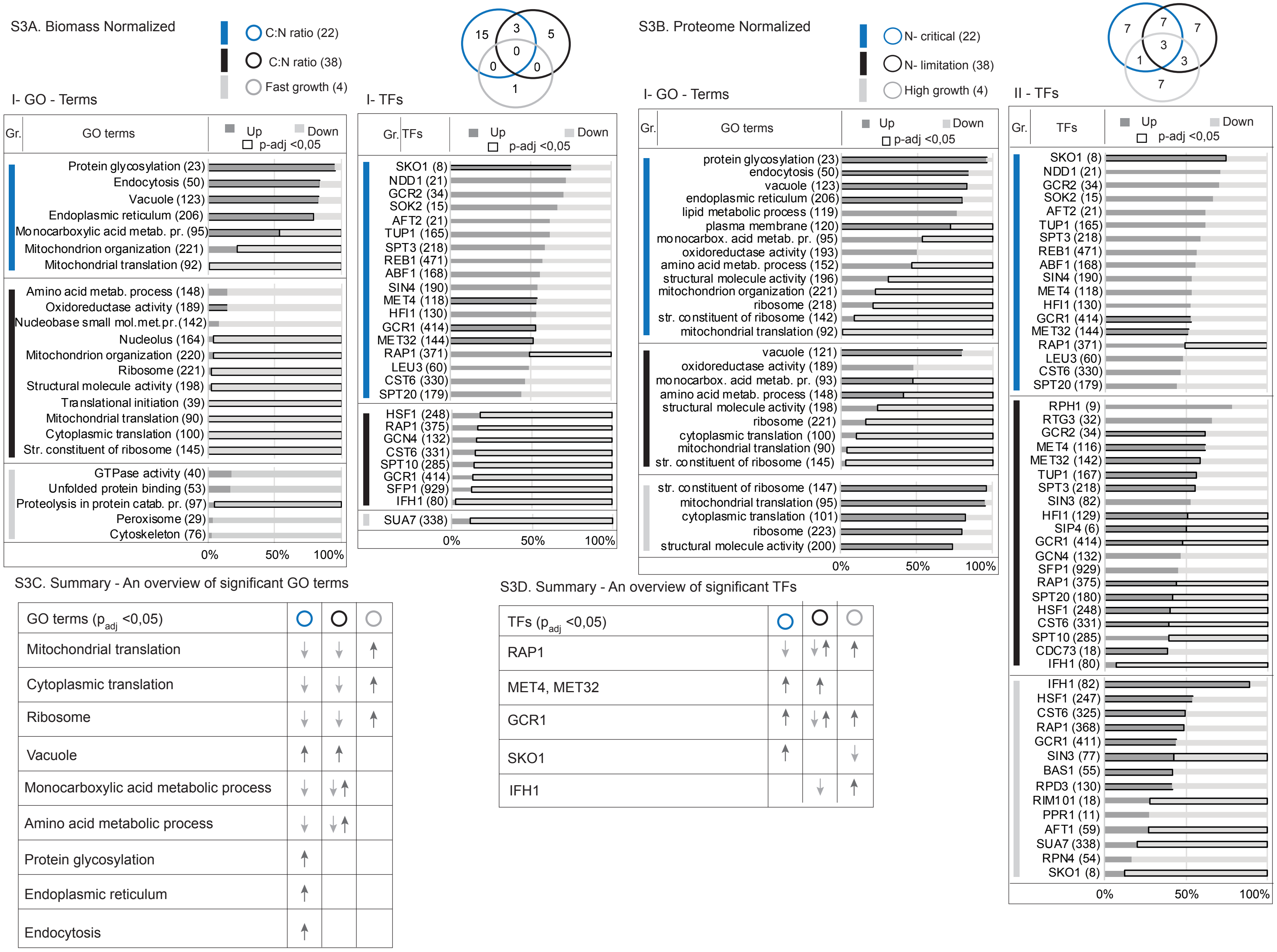

### S4A_Metabolic Pathways

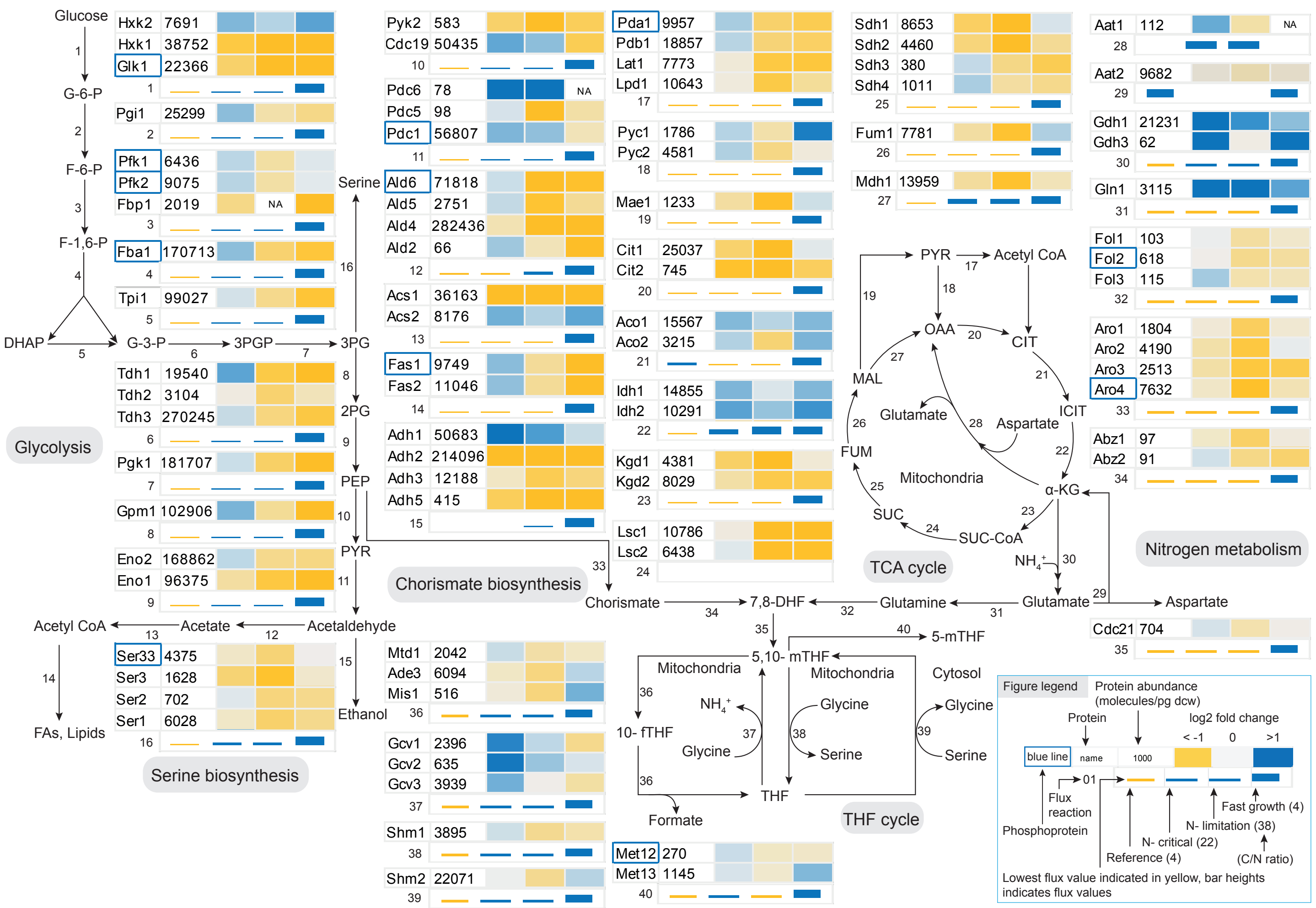
