## Supplemental M - Proteomics Data Analysis Method for "Proteome overabundance enables respiration but limitation onsets carbon overflow"

[•]Current Affiliation

### A Response-Decomposition Framework for Proteomics Data Analysis

#### 1. Proteomics data preprocessing

Protein abundance measurements were converted to numeric values, with missing measurements represented as NaNs. Abundances were divided by the smallest nonzero abundance in the dataset and transformed to the natural-log scale. The resulting log-abundance matrix was used directly for the latent-factor bootstrap and for proDA differential-abundance analysis. For the analysis purpose protein identifiers were represented by corresponding gene names, wherever necessary, such as when using common term, 'Gene Set Enrichment Analysis' or describing differential analysis.

#### 2. Bootstrap generation and differential-abundance analysis

To estimate uncertainty while preserving covariance among genes, bootstrap replicate datasets were generated using a missing-value-aware latent-factor model. Profiles were analyzed on the log scale and residualized by condition-specific means. For each gene and condition, the mean was calculated from observed measurements only; imputed values were excluded from estimation. A two-factor residual model was then fitted to the centered profiles. For gene $g$ in sample $i$, the residualized abundance was represented as a sum of gene-specific loadings on two latent factors plus gene-specific residual noise.

$$R_{gi} = \sum_{k=1}^{2} P_{gk} z_{ki} + \varepsilon_{gi}$$

where $P_{gk}$ denotes the loading of gene $g$ on latent factor $k$, $z_{ki}$ is the latent factor score of sample $i$, and $\varepsilon_{gi}$ is gene-specific residual noise.

Protein-specific residual standard deviations were estimated after removal of the fitted latent-factor contribution. To reproduce abundance-dependent and asymmetric residual variation, standardized residuals were grouped into abundance bins and the lower and upper tails were estimated separately from the 5th and 95th percentiles. The resulting tail-scale factors were interpolated across abundance levels and applied to independent Gaussian residual draws. The asymmetric residual distribution was analytically centered so that its expected value was zero.

The final synthetic abundance profile for condition $c$ was generated as

$$Y_{gi} = \mu_{g,c} + \sum_{k=1}^{2} P_{gk} Z_{k,J_i} + \varepsilon_{gi}$$

where $J_i$ is an index sampled with replacement from the observed biological replicates belonging to condition $c$, $\mu_{g,c}$ is the fitted condition-specific mean of gene $g$, $P_{gk}$ is the loading of gene $g$ on principal component $k$, $Z_{k,J_i}$ is the observed score of the sampled replicate $J_i$ on principal component $k$, and $\varepsilon_{gi}$ is centred, abundance-dependent residual noise. Complete two-factor score vectors were resampled together with replacement from biological replicates within the same condition, retaining the empirical dependence between the latent components. The original observed/missing pattern of the sampled replicate was retained, so missingness was not independently regenerated. This procedure preserved the fitted condition-specific means, the dominant low-rank covariance structure, and the observed abundance-dependent and asymmetric residual variation.

The latent-factor model was fitted only once and reused for all bootstrap replicates. Each synthetic log-abundance matrix was analyzed independently with proDA in R through RCall.jl in Julia.[1][2][3] proDA was fitted with a condition effect, and differential abundance was evaluated for each condition relative to a prespecified reference.

#### 3. Bootstrap summary statistics and GSEA ranking

For each protein and condition, the bootstrap mean differential-abundance estimate and bootstrap standard deviation were calculated across replicates. To prevent the bootstrap distribution from producing unrealistically small uncertainty for a protein whose fitted effect is precise within the synthetic replicates, the proDA standard error was used as a lower bound on bootstrap uncertainty. For each protein, the effective standard error was therefore defined as the maximum of the bootstrap standard deviation and the corresponding proDA standard error. For decomposed components spanning multiple conditions, the largest corresponding proDA standard error was used as the uncertainty floor.

The signed bootstrap ranking statistic was calculated as the mean differential-abundance estimate divided by this effective standard error. This statistic was used for downstream gene-set ranking and is referred to here as the bootstrap ranking statistic. For direction-independent analyses, the absolute value of the ranking statistic was used.

#### 4. Differential-abundance decomposition

Condition-wise differential-abundance effects were decomposed into non-overlapping shared and condition-specific components within each bootstrap replicate. For a protein with effects in the same direction across the compared conditions, the shared component was defined as the smallest absolute effect among those conditions, with its sign retained. The condition-specific component was calculated as the residual after subtraction of the shared component from the corresponding condition effect. This decomposition exactly reconstructs the original condition-wise differential-abundance effects while separating responses common to the compared conditions from residual condition-specific responses. Component-wise bootstrap ranking statistics were then calculated using the same proDA uncertainty-floor procedure described above.

### References


[1] R Core Team. R: A Language and Environment for Statistical Computing 2025.

[2] Ahlmann-Eltze C, Anders S. proDA: Probabilistic Dropout Analysis for Identifying Differentially Abundant Proteins in Label-Free Mass Spectrometry. BioRxiv 2020. https://doi.org/10.1101/661496.

[3] Lai R, Byrne S, Bates D, Alday P, Shah VB, Bouchet-Valat M, et al. JuliaInterop/RCall.jl: v0.14.13 2026. https://doi.org/10.5281/zenodo.19500543.
